# Modelling Fragile X-Associated Neuropsychiatric Disorders in Young Inducible 90CGG Premutation Mice

**DOI:** 10.1101/2025.04.24.650447

**Authors:** Gürsel Çalışkan, Sara Enrile Lacalle, Emre Kul, Miguel del Ángel, Allison Loaiza Zambrano, Renate Hukema, Mónica Santos, Oliver Stork

## Abstract

Fragile X-associated tremor/ataxia syndrome (FXTAS) is a late-onset neurodegenerative disorder caused by a preCGG repeat expansion in the FMR1 gene. Individuals with the FMR1 premutation often exhibit neuropsychiatric symptoms before FXTAS onset, leading to the identification of fragile X-associated neuropsychiatric disorders (FXAND). Rodent models of FXTAS show motor impairments, pathological intranuclear inclusions, and heightened anxiety. However, the early onset of neuropsychiatric features and underlying mechanisms remain poorly understood.

To address the above issues, we used the doxycycline (dox)-inducible 90CGG mouse model, with transgene activation at two developmental stages: adolescence and young adulthood. Mice were evaluated in a behavioural battery to assess anxiety-like behaviour, exploration, and motor coordination and learning. Next, we conducted a combination of *ex vivo* extracellular local field potential recordings to measure synaptic physiology and oscillatory activity in the limbic system, particularly in the basolateral amygdala (BLA) and ventral hippocampus (vH) regions. Parvalbumin interneurons and intranuclear inclusions in the amygdala and hippocampus were investigated by immunofluorescence, while mass spectrometry and gene set enrichment were used to identify differentially expressed proteins molecular pathways.

Adolescent 90CGG mice displayed early-onset hyperactivity, transitioning to heightened anxiety in young adulthood, coinciding with the accumulation of intranuclear inclusions in the BLA and vH. Electrophysiological analysis revealed augmented gamma oscillations in the vH, emerging during adolescence and persisting in young adulthood. These changes correlated with a reduction in parvalbumin interneurons in these regions, and together likely contribute to enhanced BLA excitability and impaired vH plasticity. Finally, proteomic analysis of the vH revealed altered proteins linked to attention deficit hyperactivity disorder in adolescence and anxiety/depression in adulthood, aligning well with behavioural findings. Importantly, these behavioural, electrophysiological, and cellular alterations were reversible upon transgene inactivation.

This study reveals a temporal progression of CGG premutation effects on behaviour, from hyperactivity to heightened anxiety to late onset motor dysfunction. Moreover, these findings provide altered network activity in the limbic system as a putative mechanism in neuropsychiatric features of premutation carriers.

## Introduction

The fragile X messenger ribonucleoprotein 1 (*FMR1*) premutation, defined by an expansion of 55–200 CGG repeats, is the genetic basis of fragile X-associated tremor/ataxia syndrome (FXTAS, [OMIM#300623](https://omim.org/entry/300623)). FXTAS is a late-onset neurodegenerative disorder characterized by progressive motor and cognitive decline, which affects approximately 40% of men older than 50 years of age^1,2^. In women, partial protection conferred by X-chromosome inactivation results in a lower prevalence of approximately 16% in older carriers^3,4^. On the other hand, 16-20% of women with the premutation who are younger than 40 develop fragile X-associated primary ovarian insufficiency (FXPOI, OMIM#311360), a form of early onset menopause^5^. In the last decade, it has become increasingly evident that the *FMR1* premutation is associated with a range of additional neurological and mental conditions including autism, attention deficit hyperactivity disorder (ADHD), anxiety, and depression^6–10^. This has led to the proposal of a new clinical entity, the fragile X-associated neuropsychiatric disorders (FXAND)^11^. *FMR1* premutation carriers face an elevated risk of developing neurodevelopmental and neuropsychiatric disorders across their lifespan^11^. As longitudinal epidemiological studies are still missing that could determine whether these conditions are early manifestations of FXTAS or distinct clinical entities, mouse models^12^ may provide valuable insights into the different pathologies associated with the premutation.

At the neuropathological level, intranuclear inclusions in neurons and astrocytes across the brain are the hallmark of FXTAS^13^. The formation of intranuclear inclusions predominantly in the cerebellum, and to a lesser extent in the amygdala and hippocampus have been demonstrated in an inducible mouse model of FXTAS (P90CGG) following the transgene activation at 4 weeks of age^14^. We have previously reported that this model manifested with gait ataxia and heightened anxiety phenotypes following the same transgene activation timeline^15^. Interestingly, inactivation of the premutation was sufficient to rescue the anxiety phenotype but not the motor phenotype, a finding paralleled by clearance of intranuclear inclusions in the hippocampus and amygdala but not in the cerebellum^15^. Effectively recapitulating multiple phenotypes characteristic of FXTAS, P90CGG model further served as a robust testing platform for therapeutic strategies targeting this disorder^16,17^. However, the physiological effects of the CGG premutation expression in the limbic areas remained unexplored. Moreover, the typical transgene induction timeline employed in prior studies that lasts 12 weeks initiating at 4 weeks of age, excludes the pre- and neonatal developmental periods, a critical window associated with emergence of a broad spectrum of neuropsychiatric conditions^18^. In this study, we sought to explore the effects of *FMR1* premutation expression in P90CGG mice during early developmental stage, a critical time window that was previously disregarded in studies using this model yet strongly implicated in the onset of neuropsychiatric phenotypes.

Of note, a preCGG *knock-in* (KI) mice FXTAS model has previously shown impaired spatiotemporal performance along with reduced hippocampal long-term potentiation (LTP) and long-term depression (LTD) induction^19^. Moreover, young preCGG KI mice exhibit robust metabotropic glutamate receptor (mGluR)-dependent LTD, which is dependent on new protein synthesis^20^. Together, evidence suggests synaptic physiology as a process worth further exploring in the context of premutation-associated neuropsychiatric phenotype.

RNA gain-of-function, characterized by the sequestration of nuclear proteins by expanded CGG repeat mRNA, and repeat-associated non-AUG (RAN) translation, resulting in the synthesis of the toxic polyglycine peptide FMRpolyG, have been proposed as key mechanisms underlying the onset and progression of FXTAS^21–23^. Numerous molecular pathways have been proposed as mediators of the neurodegeneration seen in FXTAS, including depletion of RNA-binding proteins, deficits in microRNA processing, and overburdening of the ubiquitin-proteasome system^24–26^. The pathophysiological mechanisms underlying neurodegeneration in the *FMR1* premutation are also implicated in neuropsychiatric manifestations in the absence of overt neurodegeneration^6,27^. However, the molecular pathways involved may differ and need to be identified.

In this work, we found that the activation of the premutation during early development leads to hyperactivity and heightened anxiety, but not motor dysfunction. Our behavioural findings are accompanied by the accumulation of intranuclear inclusions in the hippocampus and amygdala. Furthermore, we investigated potential physiological alterations in the amygdala and hippocampus including neuronal excitability, synaptic plasticity and network activity patterns, together with related changes in configuration of the parvalbumin interneuron population, and identified the ventral hippocampus (vH) as a putatively critical site for the observed psychopathology. Finally, we began to define molecular characteristics related to the FXAND mouse phenotype by proteomic analysis and identified a network of anxiety, hyperactivity- and depression-related factors dysregulated in the vH of the FXAND mouse model.

## Materials and methods

### Animals

Two transgenic lines were kept as independent mouse colonies: PrP-rtTA monogenic driver line and TRE-90CGG-eGFP monogenic target line. These animal lines were developed through an untargeted gene insertion method, resulting in the creation of a Tet-On system^14^. PrP-rtTA driver line includes the tetracycline-controlled transactivator (rtTA) that operates under a prion protein promotor (PrP), and TRE-90CGG-eGFP line features a tetracycline response element (TRE) upstream of a 90CGG-repeat stretch and eGFP reporter. Inducible transgenic animals were obtained by crossing female transgenic mutants carrying a TRE-90CGG repeat tract with the male transgenic PrP-rtTa driver line. At 4 weeks of age, bigenic offsprings were weaned and group housed with littermates in standard laboratory cages. Animals were kept in an inverted 12 h light/12 h dark cycle with controlled temperature and humidity conditions (lights on at 7 p.m.; room temperature 22 ± 2°C), and water and food *ad libitum*. Genotyping was performed with DNA extracted from ear punches. All animal experimental procedures were approved under local ethics committee (CEEA #42502-2-1219) and met the guidelines of the local and European regulations (European Union directive no. 2010/63/EU).

### Doxycycline induction schedule

The transgene induction started at conception and lasted a total period of 7 weeks or 12 weeks via administration of doxycycline at a concentration of 4600 mg/kg in food pellets to the breeding pairs. For experiments described below, two experimental groups were used 7 weeks after DOX+/- induction starting from embryonic development (∼4 weeks-old adolescent mice; DOX+.7w vs. DOX-.7w) and 12 weeks after DOX+/- treatment starting from embryonic development (∼9 weeks-old young adult mice; DOX+.12w vs. DOX-.12w). We also included a third group of animals that went through 12 weeks of wash-out (WO) period following the 12 weeks DOX+/- treatment (∼21 weeks old; DOX+.12w.WO vs. DOX-.12w.WO). Male P90CGG mice induced with DOX (DOX+) and their controls without DOX (DOX-) were used for behavioural, electrophysiological, immunohistochemistry and proteomic analysis. Animals were randomly assigned to each group. Animals were randomly assigned to each group, and the experimenters were blinded to the treatments to ensure unbiased results. While no formal sample size calculation was conducted, the number of animals used in each group aligns closely with those in our previous studies^15,16^.

### Behavioural tests

Motor performance in mice was assessed using the Rotarod test while anxiety like behaviour and activity were assessed using Light-dark test and elevated plus maze test. For details see supplementary material.

### Immunohistochemistry

For details on immunohistochemical methodology for assessing the density of parvalbumin-positive (PV+) interneurons and percentage of cells with FMRpolyG+ intranuclear inclusions see supplementary material.

### Electrophysiology

Acute horizontal brain slices including either ventral-to-middle hippocampus and/or lateral amygdala (LA) were prepared, and field potential recordings were performed to measure synaptic transmission, plasticity and gamma oscillations as detailed in supplementary material.

### Proteomics

#### Mass spectrometry

Mass spectrometry was performed on vH samples collected from ∼4- and ∼9-weeks-old mice with or without DOX treatment. See supplementary materials for details.

#### Statistical analyses

##### Behaviour, immunohistochemistry and electrophysiology

For the data presentation and statistical analysis of behavioural, immunohistochemical, and electrophysiological data, SigmaPlot (for Windows Version 11.0, 2008) and GraphPad Prism (Version 9, SD, California) were used. For statistical comparison of two groups, data were evaluated by Student’s *t* test (unpaired) or Mann-Whitney U test after performing normality test (Shapiro-Wilk Test) and equal variance test. Statistical analysis of behavioural rotarod data, normalized LTP data, I/O curves and PP responses were performed using repeated measures two-way ANOVA followed by *posthoc* comparison using Holm-Sidak’s multiple comparisons test with Greenhouse–Geisser correction (see corresponding figure legends for details). Grubbś outlier test was used to determine a single outlier in behavioural data. Changes in the propensity for recurrent epileptiform discharges (REDs) after CBh perfusion was statistically compared using Fisheŕs Exact test. All data are presented as mean +/- standard error of mean (SEM).

##### Proteomics

Protein identification, data analysis, and statistical validation of differential expression were performed using the Limma package in R^28^. Proteins were considered differentially expressed if they had a log fold change with a false discovery rate (FDR) less than 0.05. Gene set enrichment analysis (GSEA) was conducted using the GSEA app from the Broad Institute, employing the Molecular Signature Database (MSigDB)^29,30^. Enrichment was considered significant if the FDR was less than 0.25 and the adjusted p-value was smaller than 0.005. Identified protein hits were matched against the terms Acute Stress Disorder, ADHD, Depressive Disorder, and Anxiety from the DisGeNET database using Cytoscape for network analysis^31^.

## Results

### Anxiety-like behaviour precedes motor symptoms in the young adult FXAND mice (P90CGG.DOX+.12w)

In our previous study, we demonstrated that 12 weeks of DOX-induced P90CGG transgene activation starting at 4 weeks of age leads to motor symptoms reminiscent of FXTAS and anxiety-like phenotype^15^. In the current study, we implemented an early DOX induction schedule starting from embryonic development and continuing for 12 weeks (young adult FXAND; Fig. 1A). This allowed us to test the animals at an earlier age (∼9 weeks-old) with the same duration of DOX-induced transgene activation as in our previous study (DOX+.12w; young adult FXAND mice). Moreover, we assessed the presence of intranuclear inclusions in the lobule X of the cerebellum that was previously shown to be the most impacted region in the P90CGG model^15^ and in limbic structures related to anxiety-like behaviour, specifically the basolateral amygdala (BLA) and ventral hippocampus (vH). We identified a significant accumulation of intranuclear inclusions in the cerebellar lobule X, and substantial levels also present in the BLA, and in DG, CA3 and CA1 sub-regions of the vH (Fig. 1B). Importantly, at this stage, no obvious motor deficits were observed, illustrated by similar performance between young adult FXAND (DOX+.12w) and control (DOX-.12w) mice in the rotarod training (Fig. 1C), rotarod constant speed test (Fig. 1D) and accelerated rod test (Fig. 1E). However, we did observe a profoundly increased anxiety-like behaviour in young adult FXAND mice, confirmed by two independent behavioural tests: light-dark test and elevated plus maze (Fig. 1F-I). In the light-dark test, young adult FXAND mice spend less time in the lit compartment (Fig. 1F).

**Figure 1.**
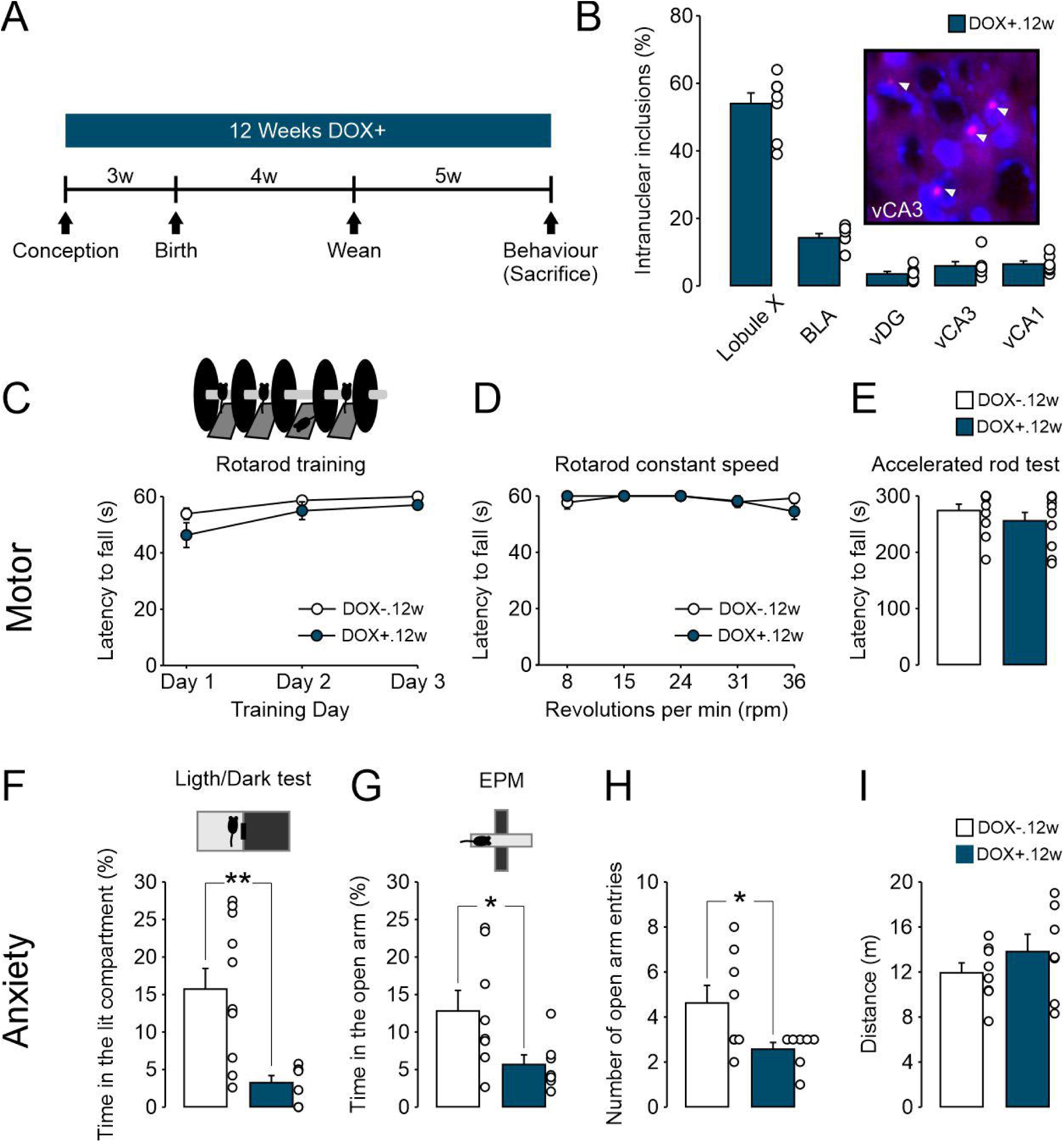
P90CGG mice treated twelve weeks with doxycycline (DOX+.12w) starting from embryonic development leads to increased anxiety-like behaviour without overt motor symptoms in young adult mice. **(A)** Experimental timeline. **(B)** Young adult P90CGG DOX+.12w mice show high inclusion load in the cerebellum lobule X (N=8 mice). Note that substantial intranuclear inclusion load is also evident in the basolateral amygdala (BLA, N=8 mice) and ventral hippocampal (vH, N=7 mice) subregions DG, CA3 and CA1. Representative intranuclear inclusions in the vCA3 of DOX+.12w adult mice are highlighted with arrows. **(C-E)** Comparable motor function between DOX+.12w (N=11 mice) and DOX-.12w (N=10 mice) mice evident by similar performance in (C) during rotarod training sessions (F(1, 19)=3.33, p=0.084), **(D)** rotarod constant speed test (F(1, 19)=0.186, p=0.671) and (E) accelerating rotarod test (U=34.5, p=0.154). **(F-I)** DOX+.12w mice show increased anxiety-like behaviour in **(F)** light-dark test (DOX-.12w: N=11 mice, DOX+.12: N=7 mice, U=8, p=0.007) and **(G-H)** elevated plus maze (EPM; DOX-.12w: N=8 mice, DOX+.12w: N=7 mice) with less time in the open arms (t(13)=2.245, p=0.043) and more number of open arm entries (t(13)=2.333, p=0.036)), with no differences in **(I)** general locomotor activity (t(13)=-1.093, p=0.294). Statistical comparison for **C, D:** repeated measures: two-way ANOVA followed by posthoc comparison using Holm-Sidak’s multiple comparisons test with Greenhouse–Geisser correction. Statistical comparison for **E, F:** Mann-Whitney Rank Sum Test. Statistical comparison for **G, H, I:** Student’s two-tailed t-test. Statistical differences are indicated via *p<0.05 and **p<0.01. Data are presented as mean ± standard error of mean (SEM). Empty circles represent individual data points.

Accordingly, in the elevated plus maze, young adult FXAND mice spend less time in the open arms (Fig. 1G) and more time in the closed arms (data not shown, t(13)=-2.405, p=0.043). Furthermore, the number of entries to the open arm were significantly lower in the young adult FXAND compared to the control mice without transgene activation (Fig. 1H). There were no changes in the total distance covered in the elevated plus maze test, suggesting comparable activity levels between control mice and young adult FXAND mice (Fig. 1I). No effects of DOX treatment were evident in control animals, which were monogenic mutant mouse lines carrying either the TRE-90CGG repeat tract or PrP-rtTA This suggests that DOX treatment by itself, without transgene activation, does not elicit anxiety-like behaviour (Supplementary Fig. 1). Of note, subsequent cessation of DOX administration for a period of 12 weeks could reduce the inclusion load in the BLA and vH (Suppl. Fig. 2A-B) and rescue the aberrant anxiety-like behaviour (Suppl. Fig. 2C-F) in young adult FXAND mice (DOX+.12w.WO). These results align well with the newly described early-onset FXAND (e.g., anxiety), which, as reported earlier^15^, can manifest concurrently with or even precede the motor symptoms linked to late-onset FXTAS^11^.

### Enhanced LA excitability and vH gamma oscillations in the young adult FXAND mice

To elucidate the potential physiological correlates that might underlie the aberrant anxiety-like behaviour in young adult FXAND mice, first, we measured synaptic transmission and plasticity in the lateral amygdala (LA) (Fig. 2A-B and Supplementary Fig. 3A). We detected an increased synaptic excitability in the LA evident by larger field excitatory postsynaptic potentials (fEPSP) in the input-output curves of young adult FXAND mice (Fig. 2A) and higher baseline transmission rates (Supplementary Fig. 3C) without any changes in the presynaptic fibre volley (FV) responses (Supplementary Fig. 3B). No changes were evident in short-term plasticity (Supplementary Fig. 3D) and long-term potentiation (LTP) (Fig. 2B).

**Figure 2.**
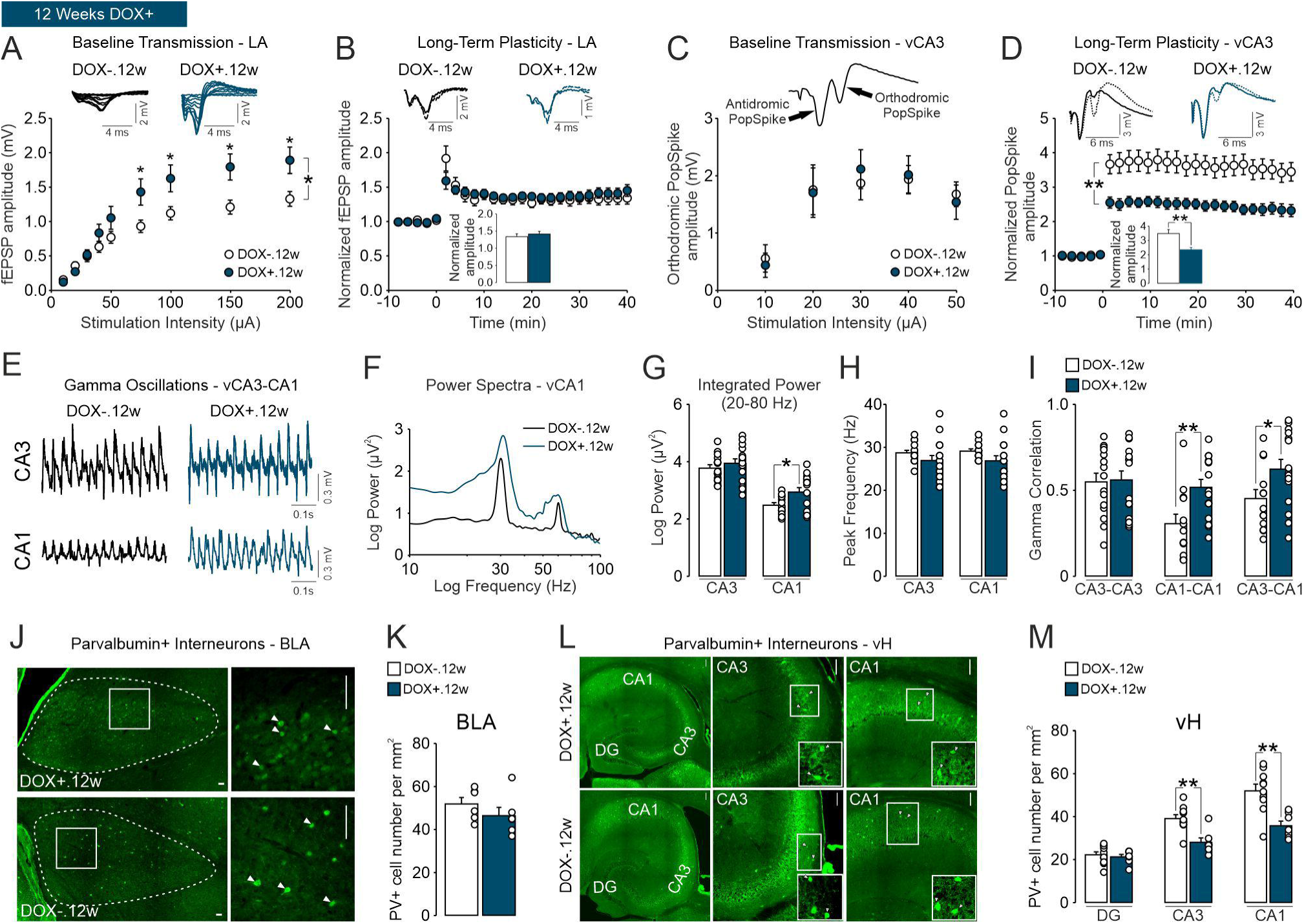
P90CGG mice treated twelve weeks with doxycycline (DOX+.12w) starting from embryonic development show aberrant enhancement of amygdalar excitability and hippocampal gamma oscillations, and a substantial reduction in parvalbumin+ interneuron number in the ventral hippocampus (vH) of young adult mice. **(A)** Comparison of input-output (I-O) curves of DOX-.12w (N=10 mice, n=31 slices) and DOX+.12w (N=7 mice, n=22 slices) mice reveal an increased baseline transmission in the lateral amygdala (LA) of DOX+.12w (F(1, 51)=5.03, p=0.029). **(B)** Similar potentiation of fEPSP responses at 30-40 min following long-term potentiation (LTP) induction (DOX-.12w: N=8 mice, n=16 slices; DOX+.12w: N=7 mice, n=12 slices, (F(1, 28)=0.303, p=0.586). **(C)** No change in synaptic transmission within the ventral CA3 (vCA3) associative network of DOX+.12w (N=5 mice, n=16 slices) in comparison to DOX-.12w (N=5 mice, n=18 slices) is evident by comparison of I-O curves for orthodromic population spikes (F(1, 32)=0.030, p=0.864). **(D)** LTP of CA3 orthodromic population spikes are significantly impaired in DOX+.12w mice (DOX-.12w: N=4 mice, n=13 slices; DOX+.12w: N=5 mice, n=13 slices, F(1, 24)=13.0, p=0.0014 and t(24)=3.358, p=0.002). **(E)** Representative traces of vCA3-CA1 and **(F)** vCA1 power spectra of cholinergic gamma oscillations induced by carbachol (5 µM) perfusion illustrating enhanced gamma oscillations in the vCA1 of DOX+.12w mice (CA3, DOX-.12w: N=5 mice, n=15 slices, DOX+12w: N=8 mice, n=16 slices; CA1: DOX-.12w: N=5 mice, n=13 slices; DOX+.12w: N=8 mice, n=16 slices). Summary data for gamma oscillations showing **(G)** increased vCA1 gamma power (20-80 Hz, CA3: t(29)=-0.864, p=0.395; CA1: t(27)=-2.397, p=0.024), **(H)** no change in gamma peak frequency (CA3: t(29)=1.345, p=0.189; CA1: t(27)=1.613, p=0.118) and **(I)** increased local CA1 correlation (CA3: U=109, p=0.901); CA1: t(26)=-2.957, p=0.007) and CA3-CA1 cross-correlation (t(26)=-2.178, p=0.039) of gamma oscillations. **(J)** Representative immunohistochemical stainings of parvalbumin (PV)+ interneurons in the basolateral amygdala (BLA) of DOX+.12w mice (N=6 mice) and DOX-.12w mice (N=6 mice) and **(K)** summary data showing no significant change in PV+ cell number in the BLA of DOX+.12w mice (t(10)=1.109, p=0.294). **(L)** Representative immunohistochemical staining of PV+ interneurons in the vH of DOX+.12w mice (N=7 mice) and DOX-.12w mice (N=10 mice) and **(M)** summary data showing substantial reduction of PV+ cell number in the vCA3 (t(15)=3.968, p=0.001) and vCA1 (t(15)=3.859, p=0.002) of DOX+.12w mice (DG: t(15)=-0.509, p=0.618). Statistical comparison for **A, B, C, D**: repeated measures two-way ANOVA followed by *posthoc* comparison using Holm-Sidak’s multiple comparisons test with Greenhouse–Geisser correction. Statistical comparison for **B** (Bar graph)**, I** (CA3-CA3 correlation): Mann-Whitney Rank Sum Test. Statistical comparison for **D** (Bar graph)**, G, H, I** (CA1-CA1 and CA3-CA1 correlation)**, K, M**: Student’s two-tailed t-test. Statistical differences are indicated via *p<0.05 and **p<0.01. Data are presented as mean ± standard error of mean (SEM). Empty circles represent individual data points.

Next, we obtained our recordings from the CA3 of vH (vCA3) and measured antidromic and orthodromic population spikes in this region (Fig. 2C-D and Suppl. Fig. 4A). We observed no changes in the magnitude of orthodromic (Fig. 2C) and antidromic (Supplementary Fig. 4B) population spikes, but a strong reduction in LTP (Fig. 2D) and short-term potentiation (Supplementary Fig. 4C). To further elucidate whether these electrophysiological changes are associated with altered network oscillations that has been linked to anxiety-like behaviour^32–34^, we measured carbachol-induced cholinergic gamma oscillations in the vCA3-CA1 (Fig. 2E-I). We detected a significant increase in the power of vCA1 gamma oscillations (Fig. 2F-G), local CA1-CA1 correlation (Fig. 2I), and CA3-CA1 cross-correlation (Fig. 2I) of gamma oscillations in young adult FXAND mice. No changes were evident in CA3 gamma power (Fig. 2G), gamma peak frequencies (Fig. 2H) and local CA3-CA3 gamma correlation (Fig. 2I).

Twelve weeks of DOX washout period (DOX+.12w.WO) was sufficient to normalize the enhanced excitability in the LA (Supplementary Fig. 5), impaired LTP in the vCA3 (Supplementary Fig. 6) and aberrant gamma oscillations in the vH of young adult FXAND mice (Supplementary Fig. 7). These results suggest that DOX-induced transgene activation and increased inclusion load in the limbic regions BLA and vH leads to enhanced LA excitability and vH gamma oscillations, which might underlie the increased anxiety-like behaviour observed in DOX+.12w mice.

### Reduced PV+ interneuron density in the vH of young adult FXAND mice

Parvalbumin-positive (PV+) interneurons are indispensable for generation of gamma oscillations^35,36^ and altered PV+ interneuron function or density in the BLA or vH have been linked to aberrant anxiety-like behaviour^34,37^. Thus, we assessed the density of PV+ interneurons in the BLA and vH of young adult FXAND mice (Fig. 2J-M). In the BLA, we observed no significant change in PV+ cell number (Fig. 2J-K). Interestingly, a significant reduction in PV+ cell density was observed in the vCA3 and vCA1 (Fig. 2L-M). However, PV+ cell density in the vDG of DOX+.12w mice was comparable to control DOX-.12w mice (Fig. 2L-M). Of note, these changes could be normalized after twelve weeks of DOX washout period (DOX+.12w.WO) (Supplementary Fig. 8). Together, these results support the notion that aberrant vH gamma oscillations and reduced PV+ interneuron density are linked to the increased anxiety-like behaviour observed in young adult FXAND mice.

### A differential behavioural phenotype is shown by adolescent FXAND mice (P90CGG.DOX+.7w)

To investigate whether the enhanced anxiety-like behaviour is already present at an earlier age, we assessed the anxiety-like behaviour in ∼4-week-old adolescent mice, which were supplied with DOX for 7 weeks starting from embryonic development (DOX+.7w; adolescent FXAND mice) (Fig. 3A). At this early stage, intranuclear inclusions were already present in the BLA and vH (Fig. 3B), though to a much lesser degree in comparison to the young adult FXAND mice (Fig. 1B). Adolescent FXAND mice had similar anxiety-like behaviour in the light-dark test and elevated plus maze to control DOX-.7w mice (Fig. 3C-E). In the light-dark test, adolescent FXAND mice spend similar time in the lit compartment (Fig. 3C). Accordingly, in the elevated plus maze, DOX+.7w mice spend comparable time in the open arms (Fig. 3D) and the closed arms (data not shown, U=30, p=0.149) as well as show similar number of open arm entries in comparison to control mice (Fig. 3E). Interestingly, adolescent FXAND mice showed signs of hyperactivity that was evident by the increased total distance covered in the elevated plus maze test in comparison to control mice (Fig. 3F). These results indicate that hyperactivity, without aberrant anxiety-like behaviour, is present from adolescence pointing to potential differences in the BLA and vH physiology and PV+ interneuron density compared to young adult FXAND.

**Figure 3.**
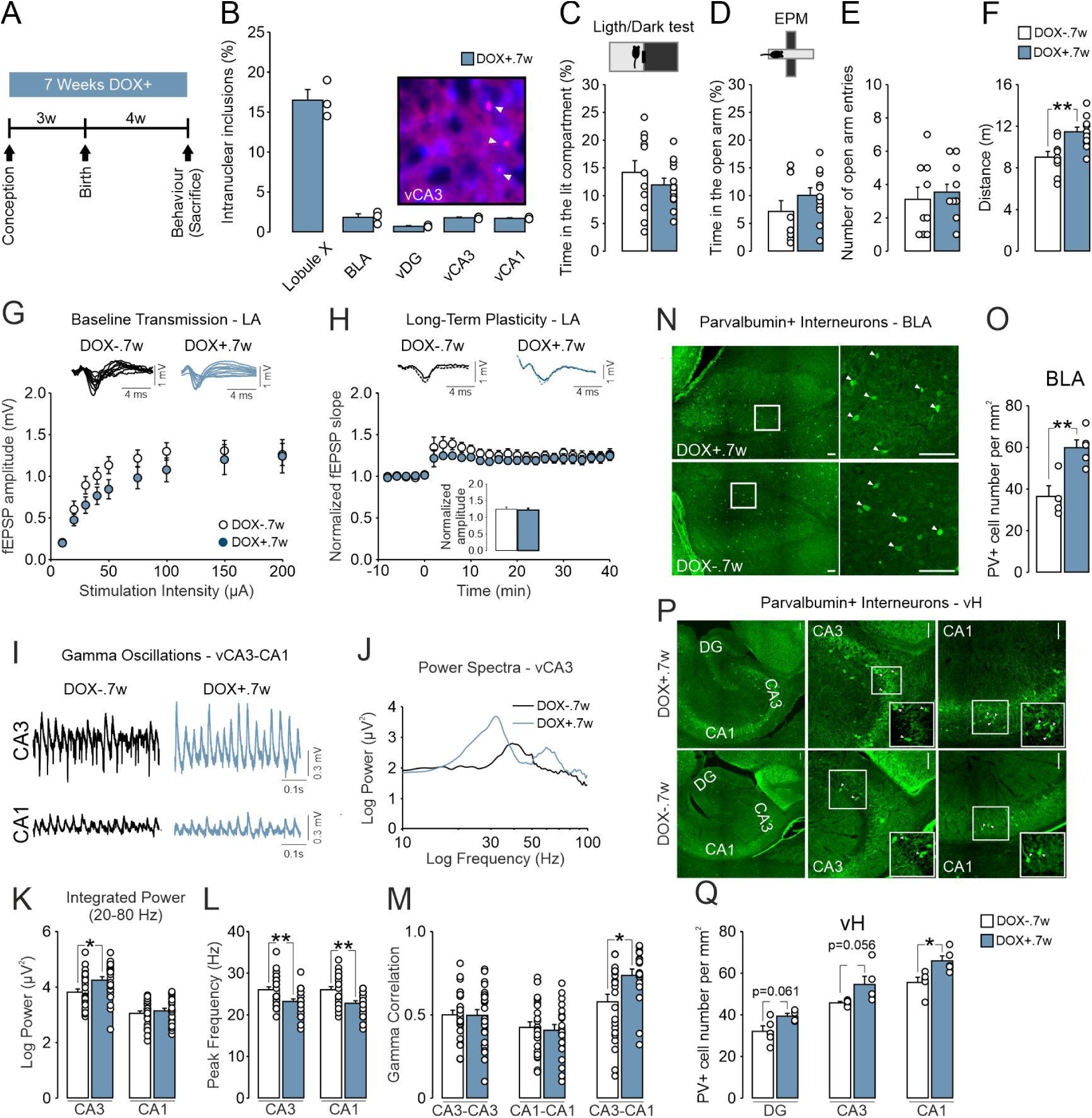
Behavioural, electrophysiological and PV+ interneuron alterations are already present in adolescent P90CGG mice treated with doxycycline for seven weeks (DOX+.7w) starting from embryonic development. **(A)** Experimental timeline. (**B)** DOX+.7w mice show moderate inclusion load in the cerebellum (N=3 mice). Note that intranuclear inclusions are also present in the basolateral amygdala (BLA, N=3 mice) and ventral hippocampal (vH, N=3 mice) DG, CA3 and CA1 subregions. Representative intranuclear inclusions in the vCA3 of DOX+.7w mice are highlighted with arrows. **(C-F)** DOX+.7w mice show unaltered anxiety-like behaviour in **C** light-dark test (DOX-.12w: N=11 mice, DOX+.12: N=12 mice, U=52, p=0.406)) and (**D-E)** elevated plus maze (EPM; DOX-.12w: N=8 mice, DOX+.12: N= 7 mice, **D:** t(18)=-1.246, p=0.229; E: t(18)=-0.514, p=0.613), but (**F)** increased locomotion in the EPM test evident by increased total distance travelled (t(18)=-3.465, p=0.003). **(G)** Input-output (I-O) curves of DOX-.7w (N=10 mice, n=23 slices) and DOX+.7w (N=10 mice, n=19 slices) mice show no alterations in baseline transmission (F(1, 40)=1.72, p=0.197) and **(H)** similar potentiation of fEPSP responses (F(1, 17)=0.861, p=0.367) at 30-40 min following long-term potentiation (LTP) induction in the lateral amygdala (LA) of DOX+.7w (DOX-.7w: N=7mice, n=9 slices; DOX+.7w: N=7 mice, n=10 slices). **(I)** Representative traces of vCA3-CA1 and **(J)** vCA3 power spectra of cholinergic gamma oscillations induced by carbachol (5 µM) perfusion illustrating enhanced gamma oscillations in the vCA3 of DOX+.7w mice (CA3, DOX-.7w: N=12 mice, n=29 slices, DOX+7w: N=12 mice, n=26 slices; CA1: DOX-.7w: N=12 mice, n=27 slices; DOX+.7w: N=12 mice, n=26 slices). Summary data for gamma oscillations showing **(K)** increased vCA3 gamma power (20-80 Hz, CA3: t(53)=-2.543, p=0.014; CA1: T(49)=-0.718, p=0.476), **(L)** reduced vCA3 and vCA1 gamma peak frequency (CA3: T(53)=3.039, p=0.004; CA1: T(48)=3.418, p=0.001) and **(M)** increased CA3-CA1 cross-correlation of gamma oscillations (CA3: t(53)=0.007, p=0.947; CA1: t(45)=0.355, p=0.724; CA3-CA1: U=118, p=0.020). **(N)** Representative immunohistochemical stainings of parvalbumin (PV)+ interneurons in the basolateral amygdala (BLA) of DOX+.7w mice (N=5 mice) and DOX-.7w mice (N=4 mice) and **(O)** summary data showing a substantial increase of PV+ cell number in the BLA of DOX+.7w mice (t(7)=-3.803, p=0.007). **(P)** Representative immunohistochemical stainings of PV+ interneurons in the vH of DOX+.7w mice (N=5 mice per region) and DOX-.12w mice (N=4-5 mice per region) and **(Q)** summary data showing a significant increase of PV+ cell number in the vCA1 of DOX+.7w mice (vDG: t(7)=-2.232, p=0.061; vCA3: t(8)=-2.228, p=0.056; vCA1: t(8)=-2.996, p=0.017). Statistical comparison for **G, H**: repeated measures two-way ANOVA followed by *posthoc* comparison using Holm-Sidak’s multiple comparisons test with Greenhouse–Geisser correction. Statistical comparison for **C, E, H** (Bar graph)**, G** (CA3-CA1 correlation): Mann-Whitney Rank Sum Test. Statistical comparison for **D, F, K, L, Q** (CA1-CA1 and CA1-CA1 correlation)**, O, K**: Student’s two-tailed t-test. Statistical differences are indicated via *p<0.05 and **p<0.01. Data are presented as mean ± standard error of mean (SEM). Empty circles represent individual data points.

### Enhanced ventral hippocampal gamma oscillations are already present in the adolescent FXAND mice

To elucidate whether altered electrophysiological phenotype in the LA and vH of young adult FXAND mice is already evident in the adolescent FXAND mice, we first assessed synaptic physiology. Intriguingly, in the LA we observed no alterations in the presynaptic FV amplitudes (Supplementary Fig. 9A), postsynaptic fEPSP (Fig. 3G) and normalized baseline transmission rate (Supplementary Fig. 9B). Adolescent FXAND mice (DOX+.7w) and control mice (DOX-.7w) had comparable LTP curves (Fig. 3H). However, we noted an increase in paired pulse responses indicating an altered short-term plasticity in the LA of adolescent FXAND mice (Suppl. Fig. 9C).

Next, we measured cholinergic gamma oscillations in the vCA3-CA1 of adolescent FXAND mice (Fig. 3I-J). To our surprise, enhanced gamma oscillations were already present at this earlier developmental stage. We detected a significant increase in the power of vCA3 gamma oscillations (Fig. 3K) and CA3-CA1 cross-correlation (Fig. 3M) of gamma oscillations in adolescent FXAND mice. Moreover, in both vCA3 and vCA1, the peak frequency of gamma oscillations was reduced (Fig. 3L). Gamma power in the vCA1 (Fig. 3K), local CA1-CA1 correlation (Fig. 3M) and CA3-CA3 correlation (Fig. 3M) of gamma oscillations remained unaltered. Of note, we have observed that a significant portion of slices exhibited recurrent epileptiform discharges upon carbachol perfusion in adolescent FXAND mice (∼38%, 16 out of 42 slices) in comparison to control mice (∼12%, 4 out of 33 slices). Such increased propensity for generation of hippocampal epileptiform activities was not present in young adult stage and following twelve weeks of DOX washout period (Suppl. Fig. 10). These results point to differential time-dependent impact of transgene activation on BLA and vH circuits and emergent network activities that are potentially linked to aberrant developmental configuration of PV+ interneurons.

### Enhanced PV+ interneuron density in the vH and BLA of adolescent FXAND mice

We also analysed PV+ interneuron densities in the BLA and vH of adolescent FXAND mice (Fig. 3N-Q). Intriguingly, in contrast to young adult FXAND mice, the adolescent FXAND mice exhibited a significant increase in the number of PV+ cells in the BLA compared to the control mice (Fig. 3N-O). A similar pattern of increased PV+ cells was observed in the vH, with this effect reaching statistical significance specifically in the vCA1 (Fig. 3P-Q). Together, these results indicate that transgene activation leads to aberrant PV+ interneuron configuration in the amygdalar and hippocampal regions that are linked to differential pathophysiological alterations across development in these limbic structures.

### Altered proteomic profile in the vH of adolescent and young adult mice

To investigate the potential molecular mechanisms underlying the anxiety and hyperactivity phenotype in FXAND mice, we performed mass spectrometry to assess protein expression changes in the vH of young adult and adolescent FXAND mice (Fig. 4a). In the adolescent mice (DOX+.7w), activation of the transgene resulted in differential expression of 45 proteins, with 33 showing upregulation (e.g., TUBA8, PFKL, HAP1, MEAF6, SLC5a3, RAB26, AP2M1, CYB5R1, PRKCA, RNH1, MTCL1, …) and 12 exhibiting downregulation (e.g., CNN2, FTL1, AHNAK, RASGRP2, LMNB1, LPP, PLXND1, LAMC1, LSS, UGT8, …). In young adult mice (DOX+.12w) bearing transgene activation, 33 differentially expressed proteins were detected, with 16 showing upregulation (e.g., IGSF21, PLCXD2, RCAN2, SDK2, IPCEF1, SYT6, CPNE5, CAMK2G, VSNL1, VAT1L) and 17 showing downregulation (e.g., PTGIS, PLIN4, ICAM1, MYH9, MYO1D, MYH10, MYH14, TESC, CNBP, NEUROD2, ZBTB20). These findings indicate that transgene activation induces time-dependent differential protein expression patterns in the vH (See Supplementary Table 1 for full list of DE proteins).

**Figure 4.**
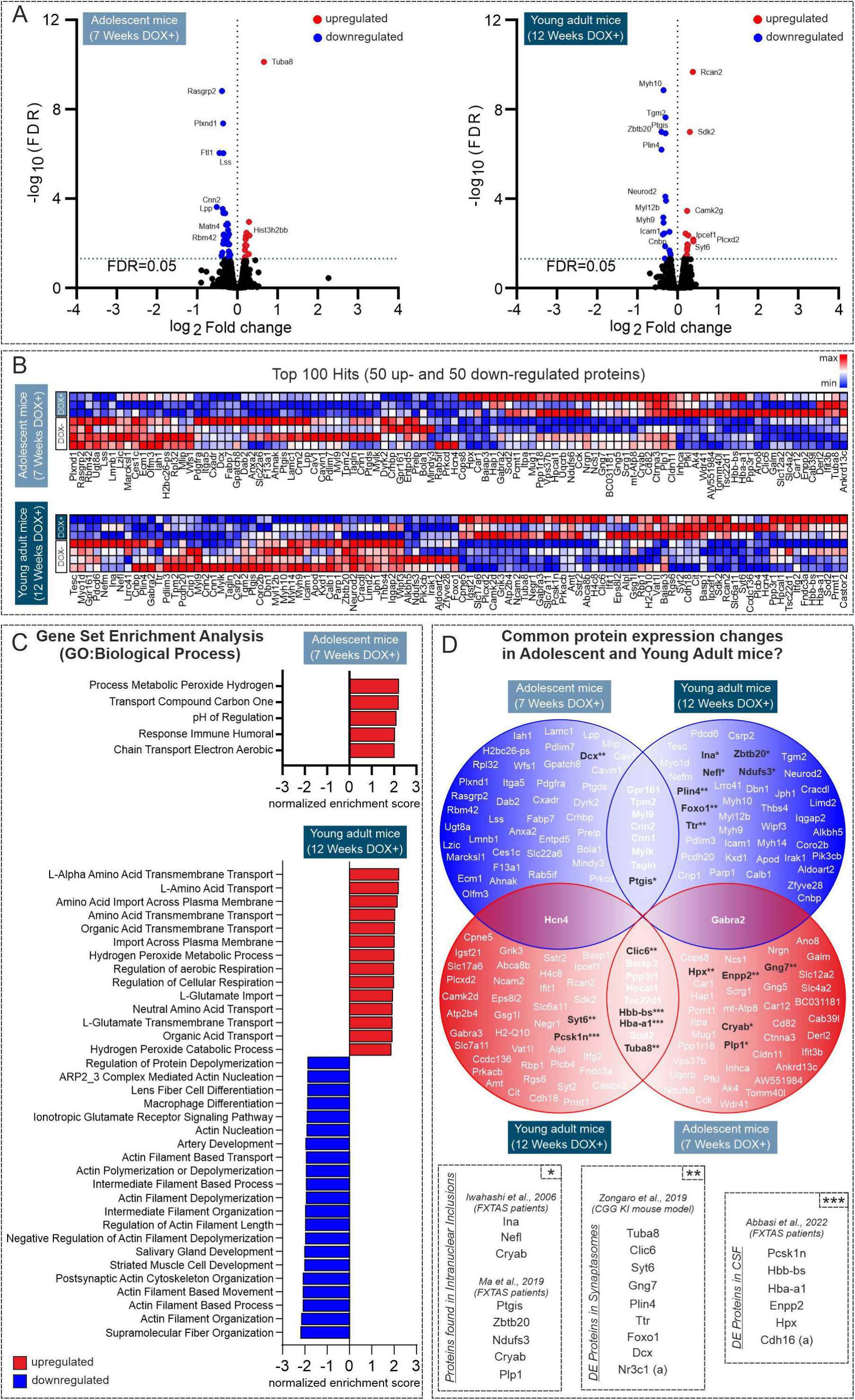
Activation of the 90CGG transgene induces time-dependent alterations in the ventral hippocampal (vH) proteome. **(A)** Mass spectrometry shows that transgene activation in adolescent (DOX+.7w) and young adult mice (DOX+.12w) alters vH proteome. The volcano plot illustrates the log2 fold change in protein expression *versus* the -log10 of the false discovery rate (FDR) for upregulated (red) and downregulated (blue) proteins with an FDR > 0.05, comparing DOX-.7w (N=4) to DOX+.7w (N=3), and DOX-.12w (N=4) to DOX+.12w (N=3). **(B)** A heatmap displays the top 50 enriched proteins in the dataset, identified through Gene Set Enrichment Analysis (GSEA) for each experimental group. The colour gradient represents expression levels, ranging from lowest (blue) to highest (red). The data were normalized row-wise using z-score normalization. **(C)** Downregulated (blue) and upregulated (red) pathways in the vH proteome of adolescent mice (DOX+.7w) and young adult mice (DOX+.12w). Bars represent the normalized enrichment score with FDR > 25% and a q-value > 0.05, based on the gene ontology (GO) biological process database. In adolescent mice, pathways related to cellular respiration, metabolism, and immune response were enriched. In young adult mice, pathways related to cellular/aerobic respiration, oxidative stress and amino acid transport were upregulated, while pathways linked to the cytoskeleton organization (e.g., intermediate filament, actin cytoskeleton) and ionotropic glutamatergic signaling were downregulated. **(D)** While adolescent and young adult mice with premutation show mainly diverging proteomic changes in the vH, several proteins with similar alterations are identified. Only two proteins (Gabra2, Hcn4) show opposite trends in their expression. Venn diagram (blue: decreased expression, red: increase expression) is based on data shown in panel b. Comparison of these data set with previously published work revealed several proteins (highlighted in black) that had been identified in the intranuclear inclusions of FXTAS patients (marked with *; ^26,38^), differentially expressed (DE) proteins in the synaptosome of CGG KI mouse model (marked with **; ^39^) and DE proteins in the cerebrospinal fluid (CSF) of FXTAS patients (marked with ***; ^40^).

Of note, most of the significant expression changes detected were moderate (∼30%). Thus, we conducted a gene set enrichment analysis (GSEA) to explore the signaling pathways and processes activated in the vH of the mice following transgene activation. This analysis revealed a group of proteins (Fig. 4b) whose differential expression affected pathways derived from the gene ontology (GO) terms related to biological processes (BP) (See Supplementary Table 2 and Supplementary Fig. 11 for complete GSEA analysis including the GO terms Molecular Function and Cellular Component). In the adolescent FXAND mice, we observed an upregulation in a limited number of pathways, including hydrogen metabolic processes, one-carbon compound transport, pH regulation, immune humoral response, and the aerobic electron chain (Fig. 4c). There was no statistically significant downregulation in any pathway (FDR<0.25). Notably, in young adult FXAND mice more pronounced changes were detected (Fig. 4c). We observed significant upregulation in pathways related to amino acid transport and metabolism (e.g., L-Alpha Amino Acid Transmembrane Transport, L-Amino Acid Transport, Amino Acid Transmembrane Transport, L-Glutamate Transmembrane Transport, Neutral Amino Acid Transport), as well as cellular respiration/oxidative stress (e.g., Regulation of Cellular Respiration, Regulation of Aerobic Respiration, Hydrogen Peroxide Metabolic Process). Conversely, cytoskeleton organization (e.g., Actin Filament Organization, Postsynaptic Actin Cytoskeleton Organization, Actin Filament Depolymerization, Intermediate Filament Organization) and glutamate receptor signaling (Ionotropic Glutamate Receptor Signaling Pathway). Overall, 12 weeks of DOX treatment had a more pronounced impact on protein expression profile in the vH, with increased amino acid transport (e.g., glutamate) and cellular respiration while negatively affecting cytoskeleton dynamics and glutamate receptor signaling. Next, among the top 50 upregulated and downregulated proteins in the adolescent and young adult FXAND mice (Fig. 4b), we identified 9 proteins that were upregulated (CLIC6, BAIAP3, PPP3R1, HPCAL1, HBB-BS, HBA-A1, SOD2, TUBA8) and 8 proteins that were downregulated (GPR161, TPM2, MYL9, CNN1, CNN2, MYLK, TAGLN, PTGIS) at both time points (Fig. 4d). Only 2 proteins showed opposing trends in their expression pattern (GABRA2, HCN4). STRING analysis of these identified proteins revealed no evidence for direct (physical) protein-protein interactions in this group of proteins (minimum required interaction score = 0.4 (medium confidence), active interaction source = Experiments). Next, we compared our top 50 upregulated/downregulated proteins and differential expression data set with publicly available data sets of previous publications (Fig. 4d). In our data set, we identified 7 proteins (INA, NEFL, CRYAB, PTGIS, ZBTB20, NDUFS3, CRYAB, PLP1) that were found in intranuclear inclusions of FXTAS patients^26,38^, 9 proteins (TUBA8, CLIC6, SYT6, GNG7, PLIN4, TTR, FOXO1, DCX, NR3C1) that were differentially expressed in synaptosome of CGG KI mouse model^39^ and 6 proteins (PCSK1N, HBB-BS, HBA-A1, ENPP2, HPX, CDH16) that were differentially expressed in the cerebrospinal fluid (CSF) of FXTAS patients^40^. These data indicate that, although the inclusion load is comparatively lower in the vH, dysregulation in molecular pathways linked to cytoskeletal organization and cellular respiration/oxidative stress is already evident in early life stages.

Last, we investigated whether the protein expression changes observed in our study are associated with neuropsychiatric disorders by comparing our results with the Diseases and Genetic Associations Database (DisGeNET) (Fig. 5). We focused on four primary neuropsychiatric disorder categories listed in the database: “anxiety”, “acute stress disorder”, “depressive disorder”, and “attention deficit hyperactivity disorder” (ADHD). We observed that the impact of transgene activation was time dependent. In adolescent FXAND mice, a larger number of proteins linked with ADHD (ITPA, POLG, PDLIM7, PTGDS, MLIP, PRKCD, and KCNLP) were identified compared to the young adult FXAND mice, where fewer proteins (SYT2, PRKCD, TBC1D24) were associated. Conversely, both adolescent mice and young adult mice showed prominent changes in the expression of proteins related to “depressive disorder” (DOX+.7w mice: LPP, WFS1, AHL1, CCK, POLG, PTGDS, ACTBL2, SOD2, CALB, LMNB1 and FABP7; DOX+.12w mice: GABRA3, AMT, PRMT1, SOD2, KHSRP, UBQLN2, ALPL, CIT, PARP1, TTR, ICAM1, CALB1, MYL12B, CNBP, NEFM and NEFL) or “anxiety” (DOX+.7w mice: DCX, BAIAP3, POLG, GABRA2, LTPA, CCK, AHL1, WFS1, LPP; DOX+.12w mice: GABRA2, MYH9, NEUROD2, INA, PCSK1N, BAIAP3, PARP1, CIT, ALPL, UUBLN2) with somewhat more predominant changes in the young adult FXAND mice. Together, these data suggest that 90CGG activation predominantly alters the expression of proteins associated with an ADHD-like phenotype during pre-pubertal stages (∼4-5 weeks old) and with anxiety/depression-associated phenotypes during early adulthood (∼9 weeks old), which aligns well with our behavioural results.

**Figure 5.**
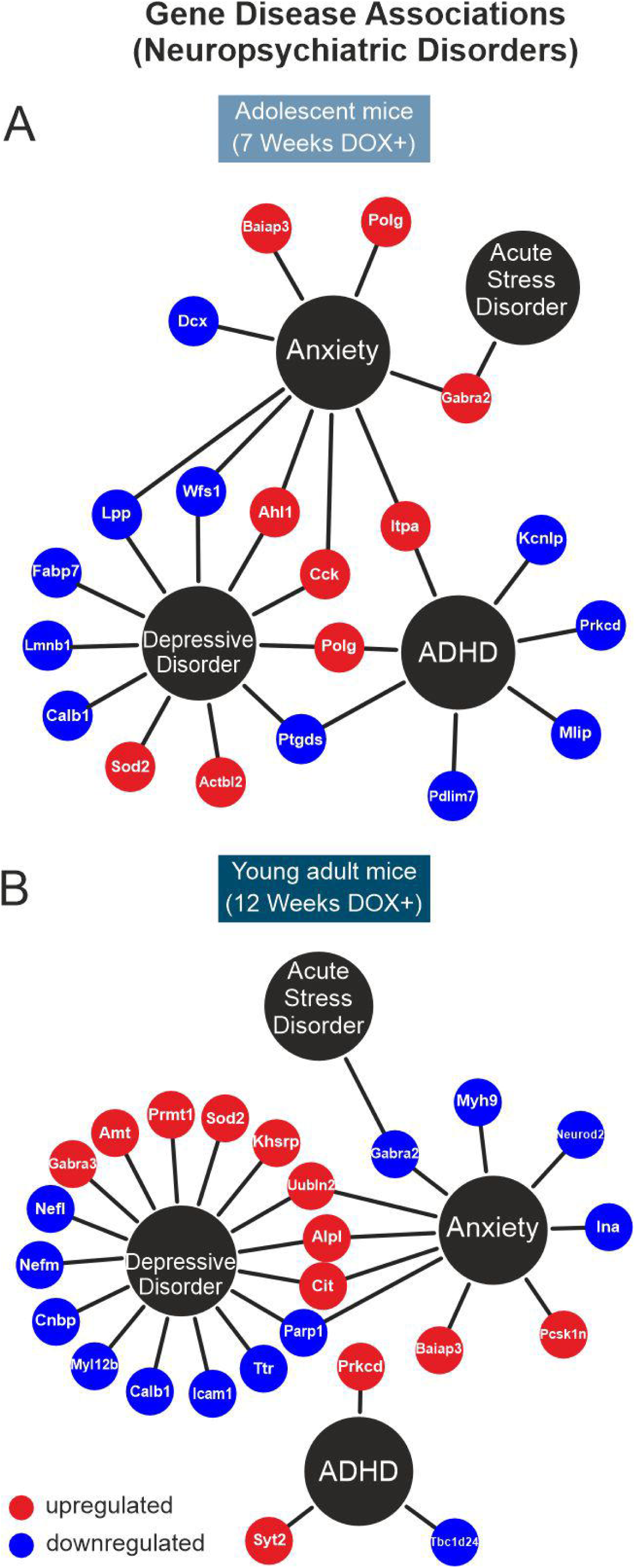
Neuropsychiatric human gene-disease associations. The top 50 downregulated (blue) and upregulated (red) proteins in (**A)** adolescent and (**B)** young adult FXAND mice with transgene activation were systematically compared to gene datasets from DisGeNet associated with Anxiety, Acute Stress Disorder, Depressive Disorder, and Attention Deficit/Hyperactivity Disorder.

## Discussion

The association of the *FMR1* premutation with neuropsychiatric features has become evident in the last years. The current study demonstrates that activation of the premutation from the embryonic stage for different durations leads to hyperactivity at adolescence and heightened anxiety in young adulthood, well before the onset of motor symptoms and thus establishes the inducible P90CGG young mouse as a model of FXAND. Furthermore, this work identifies increased vH network oscillations and enhanced BLA excitability, which could result from changes in the configuration of PV+ interneurons in these brain regions, potentially underlying the impact of the premutation on heightened anxiety levels. Finally, activation of the premutation shows a temporal effect on the molecular composition of the vH, which correlates with the progressive emergence of different neuropsychiatric disorders, including ADHD, anxiety and depressive disorder, in individuals carrying the premutation.

### Premutation activation from the embryonic stage and its contribution to FXAND

Expression of the premutation in different temporal windows results in specific phenotypes. Beyond FXTAS, the *FMR1* premutation has been associated with a range of neurodevelopmental and neuropsychiatric conditions. In particular, young boys with the premutation, especially at an early age, often exhibit mild developmental delays and show increased rates of autism spectrum disorder (ASD) and ADHD symptoms when compared to their non-carrier siblings^6,9^. Moreover, anxiety and mood disorders are among the most common psychiatric conditions affecting individuals who carry the premutation^10^. Anxiety disorders usually begin in childhood and are present during the entire lifetime of premutation carriers^8,10,11^. In line with the anxiety phenotype, premutation carriers show altered amygdala activation^27^, a key brain region in emotional processing of anxiety and fearful stimuli^41^. Here, we show that activation of the premutation starting from embryonic development leads to earlier hyperactivity followed by heightened anxiety. On the other hand, the activation of the premutation starting in young adulthood results in increased anxiety and motor impairments resembling FXTAS^15^. It remains under investigation whether these early neuropsychiatric features represent distinct clinical entities or are prodromal signs of late-onset FXTAS. However, the clearance of intranuclear inclusions from the BLA and vH, along with the simultaneous reduction in anxiety upon premutation inactivation may suggest distinct disease mechanisms.

### Impairments in synaptic plasticity and network oscillatory activity in the vH may be linked to heightened anxiety

Analysis of synaptic function in young adult and adolescent FXAND mice revealed developmental stage-dependent effects on the synaptic excitability and plasticity in the vH, a region that plays a major role in anxiety^41^. Our results confirm the deleterious effects of the premutation in synaptic function, as previously described^19,20^. In previous experiments authors showed that dorsal hippocampus (dH) slices from CGG premutation KI mice have unchanged basal synaptic function^20^, but significantly less LTP induction than wild-type slices, which could account for the observed alterations in spatiotemporal processing^19^. Likewise, in our study, the activation of the premutation in P90CGG.DOX+ animals resulted in a decreased LTP in vH slices.

The hippocampus has been recognized as a key structure in the generation of distinct neural network oscillations, which mediate neural communication and plasticity within and across limbic brain regions^32,42^. The observed alterations in synaptic plasticity may be intricately linked to alterations in brain rhythms, particularly in the hippocampus^43^. We indeed observed increased gamma-range oscillations in the vH together with increased vCA3-CA1 communication, as indicated by high synchronous activity. The identified alterations in vH oscillatory activity may be crucial for regulating the synaptic plasticity processes essential for adaptive behaviour^42^.

Brain network oscillations are a hallmark of multiple brain disorders^44,45^ and specifically, vH oscillations in the theta and gamma ranges have been associated with anxiogenic features (reviewed in^32,34,46,47^). Thus, alterations in neuronal oscillations observed in the P90CGG premutation model may comprise a mechanism through which disturbances in hippocampal function result in anxiety. The increased gamma rhythmicity could impair hippocampal functionality, exacerbating anxiety, as is seen in other neurological conditions such as epilepsy^48^. Supporting this, increased hippocampal gamma oscillation rhythmicity has consistently been correlated with heightened anxiety in humans^49–51^. In line, recent evidence from our laboratory demonstrates that *ex vivo* gamma oscillations in the vH are elevated in inbred mouse strains with higher anxiety and stress-susceptibility^52^.

The increased hippocampal gamma rhythmicity seen in the P90CGG transgenic mice induced at embryonic stage may also lead to increased activity in interconnected brain regions, such as the amygdala. In line with this, we detected an increase in excitability of amygdala as one of the potential correlates of increased in anxiety in the young adult FXTAS mice. Indeed, emergence of amygdalar theta oscillations and their increased correlation with hippocampal theta oscillations are well-established hallmarks of fear and anxiety^32,46,53^.

### Altered configuration of PV+ interneurons: potential link to aberrant gamma oscillations and anxiety

Parvalbumin-positive (PV+) inhibitory interneurons are highly vulnerable to stress and have been associated with various neuropsychiatric disorders, including anxiety, depression, and autism spectrum disorder^54^. Notably, dysfunction of PV+ interneurons appears to be a central feature in several neurodegenerative diseases characterized by comorbid cognitive and emotional impairments, such as Alzheimer’s disease and various forms of dementia^55^. Our study revealed that in the prodromal stages of the neurodegenerative condition FXTAS— spanning adolescence to young adulthood—PV interneurons initially exhibit an increased density, which is subsequently followed by a marked reduction in early adulthood. Further research is needed to clarify whether the changes in PV density reflect alterations in PV expression or an actual loss of PV interneurons.

Alterations in PV interneuron density or function are closely linked to gamma oscillations^56,57^. Numerous studies have further demonstrated the causal role of PV interneuron activity in sustaining hippocampal gamma oscillations both *in vivo* and *ex vivo*^58–60^. Specifically, PV interneuron activity in the vH plays a pivotal role in regulating anxiety-like behavior^61,62^. Aligning well with the current data, in our recent study, mouse strains with heightened trait anxiety—measured using tasks similar to those employed here—exhibited increased *ex vivo* gamma oscillations in the vH, along with changes in PV interneuron density along the vCA3-CA1 axis^52^. Moreover, elevated anxiety or emotionality appears to enhance *in vivo* gamma oscillations in the vH, with hypersynchronous gamma activity being linked to anxiety-related neuropsychiatric disorders^33,49,63^. Of note, reduced PV interneuron number and increased *ex vivo* and *in vivo* hippocampal gamma oscillations have been also observed in *Fmr1* KO mice^64–67^. Thus, early-life alterations in PV interneuron configuration may profoundly affect behaviourally relevant network activities, such as gamma oscillations. These disruptions could contribute to the hyperactivity observed in adolescent FXAND mice and the increased anxiety seen in young adults.

Oxidative stress has been shown to impair the function and survival of PV interneurons, which are particularly vulnerable to metabolic and oxidative insults due to their high energy demands^68^. Previous studies have reported altered expression of proteins involved in mitochondrial bioenergetics and oxidative stress in FXTAS patients^26,38^. Using a transgenic mouse model, our study now demonstrates that even during the earlier life stages (adolescence to young adulthood), in the prodromal phase of FXTAS, there is an upregulation of molecular pathways related to cellular respiration and oxidative stress. Notably, a previous study has shown that the ventral portion of the hippocampus is particularly susceptible to oxidative stress, leading to the loss of PV interneurons, disrupted gamma oscillations, and anxiety-like behaviour^69^. This suggests that elevated metabolic demands during these early stages may contribute to PV dysfunction and/or loss, potentially initiating a cascade of events that disrupt gamma oscillations (Fig. 2-3) and increase susceptibility to epileptiform activity (Suppl. Fig. 10).

Accumulating evidence suggests that impairments in GABAergic signaling, which is critical for the generation of gamma oscillations and the regulation of anxiety-like behaviour, play a key role in FXTAS pathophysiology. In KI CGG mice harbouring the premutation, GABA signaling was found to be upregulated in the cerebellum but not in the cortex^70^. Additionally, cultured hippocampal neurons from KI CGG mice exhibit cluster burst firing activity associated with GABA deficits, a phenotype that can be ameliorated by treatment with allopregnanolone, a GABA_A_ receptor agonist^71^. Furthermore, allopregnanolone treatment has been shown to effectively reduce anxiety symptoms and to restore GABA metabolism in individuals with FXTAS^72,73^. Consistent with these findings, our data reveal deficits in readouts closely linked to GABAergic signaling in the vH of P90CGG mice.

### 90CGG activation has a time-dependent impact on the molecular composition of vH

Both in humans and in mice, the premutation enhances *Fmr1* transcription, while impairing protein translation^12,21,74^. The *FMR1* premutation is hypothesized to elicit a gain-of-function toxicity at the mRNA level, either by sequestering RNA-binding proteins^75^, or promoting repeat-associated non-AUG (RAN) translation^22,23^. Although the 90CGG tract of the P90CGG mouse model is expressed outside the context of the *Fmr1* gene, it enables the investigation of both pathogenic mechanisms and their contributions to observed phenotypes^14^. Our previous^15^ and current data support the role of RAN translation toxicity in FXTAS and FXAND phenotypes, mediated by polyG aggregate formation in brain regions linked to these phenotypes. However, the effects may be limited to specific behavioural domains, as no correlation was observed between the presence of polyG-positive aggregates and cognitive behaviour in the forebrain-specific inducible mouse model expressing 103CGGs under the control of the CamKIIa promoter^76^.

To investigate the pathogenic molecular alterations caused by the CGG premutation, we performed a proteomic analysis in the vH of P90CGG mice treated with DOX for 7 weeks and 12 weeks. First, the data show a time-dependent impact of the premutation activation on the molecular profile of the vH. Second, although the inclusion load is comparatively lower in the vH, dysregulation in molecular pathways linked to cytoskeletal organization and cellular respiration/oxidative stress is already evident in early life stages. Thus, these dysregulated pathways may already contribute to disease pathogenesis associated with FXAND. This finding aligns with previous research, which also showed increased levels of proteins such as α/β-crystallin (CRYAB), intermediate filament proteins, microtubule-associated proteins, myelin-associated proteins, and proteins related to mitochondrial bioenergetics and oxidative stress (Figure 4)^26,38^. To determine whether the identified proteins are associated with neuropsychiatric conditions relevant to FXAND, we conducted a bioinformatic analysis of protein composition changes linked to four key neuropsychiatric conditions commonly observed in *FMR1* premutation carriers: ADHD, anxiety, acute stress disorder, and depressive disorder^6,8–10^. Interestingly, molecular data correlate short-term activation (∼7 w) of the premutation with ADHD, while extended activation (∼12 w) is associated with depressive disorder, mirroring the progressive development of these symptoms in humans carrying the premutation. In addition, the protein alterations observed following both 7 weeks and 12 weeks DOX induction were linked to anxiety, a symptom that typically manifests in early childhood and persists until late adulthood in individuals with the premutation^10,11^.

## Conclusion

The P90CGG model overexpresses an expanded CGG repeat tract independent of the *FMR1* gene, and its artificial design necessitates careful consideration of its translational relevance. Nonetheless, it serves as a robust *in vivo* platform that replicates a broad spectrum of premutation-associated phenotypes, many of which are absent in more natural models, such as KI models. Additionally, the model’s inducibility offers temporal control over transgene expression, enabling the study of multiple premutation-related disorders and allowing for the targeted cessation of expression, thereby facilitating the evaluation of therapeutic time windows. In the P90CGG mouse model, we demonstrate an age-dependent impact of the premutation on vH and amygdala resulting in hyperactivity and anxiety during a time that corresponds to a period earlier than the motor impairments commonly reported in FXTAS. This work further reveals a physiological mechanism that links premutation-induced aberrant gamma oscillations in the vH to the early development of anxiety-related FXAND symptoms. Activation of the premutation alters the configuration of PV+ interneurons and the molecular composition of vH, influencing synaptic function and network oscillations. This disruption might contribute to the disturbed behavioural output, particularly related to neuropsychiatric manifestation.

## Data availability

Proteomic data are available as supplementary material. Behaviour, electrophysiology and immunohistochemistry data that support the findings of this study are available from the corresponding author, upon reasonable request.

## Acknowledgements

We are particularly grateful to R. Willemsen for his support in model development and scientific discussions. We thank F. Blitz for the excellent work with the immunohistochemistry stainings, A. Koffi von Hoff and S. Stork for excellent technical assistance and to A. Bohnstedt and D. Al-Chackmakchie for excellent animal care.

## Supplementary material

Supplementary material is available at *Brain* online.

## Authors’ Contributions

GC, MS, SEL, ALZ, EK, MdA contributed to data acquisition and data analysis. GC, EK, MS, OS contributed to study design. GC and MS wrote the first draft of the manuscript with input from other authors. GC, MS, OS acquired funding and contributed to supervision of the study. All authors critically reviewed the manuscript for important intellectual content and confirmed the last version of the manuscript.

