## Supplementary Information for "Modelling Fragile X-Associated Neuropsychiatric Disorders in Young Inducible 90CGG Premutation Mice"

**Including Supplementary Figures, Materials and Methods**

### Supplementary Materials and Methods

#### Behavioural tests

##### Rotarod test

Motor performance in mice was assessed using the Rotarod test. During the training session, mice were placed on a rotating rod set to a constant speed of 15 rpm for a maximum of 60 seconds over four trials. This training occurred over one day to evaluate motor performance related to treatment strategies and was repeated for three consecutive days during the phenotyping experiments. Following training, mice were tested at various constant speeds (8, 15, 24, 31 and 36 rpm), with two trials conducted at each speed, lasting up to 60 seconds. On the final day, the mice underwent a 5-minute accelerated ramp test, where the rod's speed increased from 4 to 40 rpm across four trials. The latency to fall from the rod was recorded for each trial[1]. The apparatus was cleaned with 10% ethanol after each session.

##### Light-dark test

Anxiety-like behaviour was assessed using a chamber equipped with a motion-detecting photobeam system and a light source. The setup consisted of a chamber divided into two 20 cm by 20 cm compartments, separated by a methacrylate box. These compartments were connected by a four-by-four cm opening. One compartment remained in complete darkness (0 lux), while the other was illuminated at 110 lux. The system was cleaned with 10% ethanol and the mice were placed in the illuminated compartment and allowed to explore the entire apparatus for five minutes. The total time spent in each compartment was recorded and calculated as a percentage of the total time.

##### Elevated Plus maze

This test aims to study the anxiety-like behaviours based on a natural tendency of animals to investigate a new environment *versus* the negative properties of an elevated open hallway. Mice were placed in the centre of the system and allowed freely to explore the maze for five minutes while being video recorded and tracked by the Anymaze software. This setup has two closed arms, with walls 40 cm in height and two open arms. The maze is elevated 50 cm from the floor and the arms are 21 cm long. The percentage of time spent in the open arms was calculated in addition to the number of entries to the open arms and the total distance travelled.

#### Immunohistochemistry

#### Immunohistochemistry for Parvalbumin (PV)

After mice were anesthetized with isoflurane, the brains were removed and post-fixed with 4% paraformaldehyde (PFA) for 24 hours at 4 °C and cryoprotected with 30% sucrose in phosphate-buffered saline (PBS) until brains sank. Brains were then snap frozen using liquid-nitrogen-cooled 2-methylbutane. Free-floating 30 μm thick horizontal sections containing ventral hippocampus (vH) and the amygdala were cut on a cryostat and used for immunohistochemical experiments. Free-floating sections were washed with PBS and treated with 0.01M sodium citrate/0,05% triton X-100 (pH=6) for 30 minutes in a hot air oven at 95 °C for antigen retrieval. For permeabilization, slices were incubated with 0.3% triton X-100 in PBS for 10 minutes at room temperature (RT). Then, the sections were blocked in 4% bovine serum albumin (BSA) for 1 hour at RT. Rabbit polyclonal parvalbumin antibody (#ab11427, Abcam, RRID: AB_298032) at a dilution of 1:500 was used as a primary antibody and sections were incubated in primary antibody solution (2% BSA in PBS) overnight at 4 °C. Later, sections were washed in PBS followed by 1 hour incubation at RT with anti-rabbit secondary antibody Alexa-Fluor 488 (1:1000; #A32731, Invitrogen, RRID AB_2633280) secondary antibody in the same incubating solution as the respective primary antibody. Sections were incubated with DAPI (300 nM) for nuclear counterstaining. A Leica microscope DMi8 (20×Dry objective, LAS X software with THUNDER technology) was used to acquire the Z-stack images (30 μm stack volume with a step size of 5 μm) of the whole area of the hippocampus and basolateral amygdala (BLA). For quantification of PV+ cells in the vH, four sections per animal were stained and evaluated in the stratum radiatum, stratum pyramidale and stratum oriens of both, CA1 and CA3, and the molecular layer, granular layer and hilus of the dentate gyrus. In addition, four sections per animal were stained for analysis of the BLA. The Z-project tool in the ImageJ software (version 2.1.0/1.53c) was used to obtain the maximal projection images from each stack. All the hippocampal areas and BLA area were marked (region of interest-ROI) based on DAPI staining using the free hand function in ImageJ. Cell numbers were counted manually using the Cell Counter plugin. Overlapping cells were checked using DAPI counter staining to avoid occurrences of miscounting. Area sizes were measured, and the cell counts were reported as number of positive cells per $mm^2$. The average for each individual animal was calculated and used for statistical comparison between groups.

#### Immunohistochemistry for Ubiquitin and FMRpolyG peptide

Similar procedure as above was used for post-fixation and cryoprotection of the brains. Eight-μm sections were sliced using a cryostat and collected directly to poly-L-lysine coated slides.

Sections were treated with 0.01 M sodium citrate (pH=6.0) in a microwave oven for antigen retrieval. An extra antigen retrieval step with proteinase K (5 µg/ml) of 10 min at 37 °C was included in the protocol. Intranuclear inclusions were identified using either primary antibody mouse anti-FMRpolyG or rabbit anti-ubiquitin. For detection with mouse anti-FMRpolyG, sections were pre-treated to block mouse immunoglobulins via M.O.M. reagents (Vector) for 1 hour. They were then incubated with primary antibody mouse anti-FMRpolyG (8FM, 1:200) in primary antibody incubation solution from M.O.M reagents (Vector) at 4 °C overnight. Following this, biotinylated secondary antibody incubation was performed using M.O.M reagents (Vector) for 10 minutes, and the antigen-antibody complexes were visualized by staining with Cy5 streptavidin (Invitrogen) 1:1000 for 30 minutes at RT. Sections were counterstained with DAPI and coverslipped with Immu-Mount™ (Shandon).

For detection with rabbit anti-ubiquitin, endogenous peroxidase was blocked using a PBS solution containing 0,6% $H_2O_2$ and 0,125% sodium azide Sections were then blocked with a general protein blocking solution (PBS containing 0,5% protifar (Nutricia) and 0,15% glycine (Sigma)) and incubated overnight at 4 °C with the primary antibody rabbit anti-ubiquitin (Dako Z0458; 1:250). Antigen-antibody complexes were visualized by incubation with DAB substrate (Dako K3468) after incubation with Brightvision poly-HRP-linker (Immunologic; DPVO-HRP55). Slides were counterstained with hematoxylin and mounted with Entellan™.

For intranuclear inclusion quantification, fluorescent images of cerebellum lobule X, vH and BLA were taken. Images were captured with a z-step size of 2 µm under 40x magnification on an epifluorescence microscope (Leica) and subsequently analyzed with the open-source image processing software ImageJ. For each brain region, DAPI+ nuclei were counted (in cerebellum, 400 DAPI+ nuclei within the granular layer of the lobule X with a grid of 150 µm x 150 µm, 400 DAPI+ nuclei in the CA3 and CA1 pyramidal layers and in the DG granule layer in the vH using a grid of 150 x 150, 200 DAPI+ nuclei in the hilus using a grid of 250 x 250, and 200 DAPI+ nuclei in the BLA with a grid of 250 x 250). For this quantification random images were used that were generated from whole lobule X in cerebellum, BLA and vH images by a custom cell counter script developed for ImageJ software. For each DAPI+ nucleus counted, the script recorded whether the nucleus contained an FMRpolyG+ intranuclear inclusion. Based on the data from the identified inclusions, the percentage of FMRpolyG+ nuclei were calculated.

#### Electrophysiology

##### Slice preparation

Male mice were deeply anesthetized with isoflurane and decapitated. Brains were rapidly (~30-60 sec) removed and placed in cold (4-8 °C) carbogenated (5% $CO_2$/ 95% $O_2$) artificial cerebrospinal fluid (aCSF) containing (in mM) 129 NaCl, 21 $NaHCO_3$, 3 KCl, 1.6 $CaCl_2$, 1.8 $MgSO_4$, 1.25 $NaH_2PO_4$ and 10 glucose. Horizontal brain slices (400 µm) including either ventral-to-middle hippocampus and/or lateral amygdala (LA) were cut at an angle of about 12° in the fronto-occipital direction as reported previously[2,3]. Three to four slices per hemisphere were transferred to an interface chamber perfused with aCSF (at 32.0 ± 1.0 °C; flow rate: 2.0 ± 0.2 ml / min, pH 7.4, osmolarity ~300 mosmol / kg). Slices were incubated for at least 1 h before starting recordings. During execution of the experiments the experimenter was blind to the treatment of mice.

#### Field potential recordings

Glass electrodes filled with aCSF (~1 MΩ) were used for extracellular field potential recordings and placed at a depth of 70-100 µm in the slice tissue. Signals were pre-amplified using a custom-made amplifier and low pass-filtered at 3 kHz. Signals were sampled at a frequency of 10 kHz and stored on a computer hard disc for off-line analysis.

##### Baseline Synaptic Transmission and Plasticity

For LA electrophysiology, the recording electrode was placed in the LA and the stimulation electrode was placed at the external capsule (EC) for stimulation of cortical input to LA. To assess synaptic transmission and plasticity at the ventral CA3 associative network, the recording electrode was placed at the stratum pyramidale of the CA3b. The stimulation electrode was placed at the stratum radiatum of proximal CA1 for stimulation of Schaffer collaterals (SC) and commissural/associational fibers. This stimulation configuration triggers a fast antidromic population spike (PS) in the CA3 stratum pyramidale followed by an orthodromic PS as a result of activation of local CA3-CA3 synapses. At least 20 minutes of baseline responses were recorded (0.033 Hz, pulse duration: 100 µs) to make sure that the responses were stabilized before measurement of baseline transmission. For CA3-CA3 recordings, an input-output (I/O) curve was obtained using five intensities ranging from 10 µA to 50 µA and for EC-LA recordings an input-output (I/O) curve was obtained using nine intensities ranging from 10 µA to 200 µA. The stimulus intensity was adjusted to elicit ~40-50% of the maximum amplitude and was further used for the paired-pulse (PP) and long-term potentiation (LTP) measurements. PP responses were obtained using intervals from 10 ms to 500 ms. After a 10 minutes of baseline recordings (0.033 Hz), LTP at the CA3-CA3 auto-associative synapse and at the EC-

LA path was induced by two high frequency stimulation (HFS) trains (100 Hz, 20 s interval, 1 sec duration, 100 stimuli per HFS). After LTP induction responses were recorded for 40 minutes (0.033 Hz). Evoked fEPSPs and PSs were analyzed offline using self-written MATLAB-based analysis tools (MathWorks, Natick, MA). FV amplitude was calculated using peak to through amplitudes of descending phase of the FV. PP was calculated by dividing the second fEPSP amplitude to the first fEPSP amplitude. For EC-LA synapse, recorded negative waveform corresponds to a summation of both EPSPs and synchronized action potentials (population spike component)[4,5]. Thus, the data for EC-LA synapse is presented as amplitude to reduce the variability as seen for the slope measure in the LA[5]. For the analysis of CA3-CA3 synapse, PS amplitudes were calculated by averaging peak-to-peak amplitudes of descending and ascending phases of the PS. PP in the CA3-CA3 synapse was calculated by dividing the amplitude of the 2nd orthodromic PS to the amplitude of the 1st orthodromic PS. fEPSP and PS amplitudes obtained during LTP measurements were normalized to the average of baseline data 10 minutes before the induction of LTP. The data was further binned into two minutes data points (average of 4 values).

**Carbachol-induced gamma oscillations**

Carbachol (CBh) was purchased from TOCRIS (Bristol, UK). Stock solution including 50 mM CBh was diluted to a final concentration of 5 µM in aCSF prior to the experiment and applied via continuous bath perfusion. Before CBh perfusion, the recording chamber temperature was increased to 35 °C. After 45-60 minutes perfusion of CBh, glass electrodes were placed at the *stratum pyramidale* of area CA3 and CA1. Three-to-five minutes recordings were obtained. Gamma oscillations were analyzed using custom-made Spike2 scripts (Cambridge Electronic Design, Cambridge, UK). Power spectra were generated from two minutes records with a frequency resolution of 0.8192 Hz. Peak frequency (Hz, the frequency value at the maximum power value) and integrated power ($\mu V^2$) between 20 to 80 Hz were calculated. Slices with peak powers lower than 10 $\mu V^2$ / Hz and peak frequencies lower than 20 Hz were discarded to assure inclusion of slices with good gamma activity. Integrated power values were further converted to logarithmic scale for further analysis. For autocorrelation (CA3-CA3 or CA1-CA1) and cross-correlation (CA3-CA1) analysis data was low-pass filtered at 100 Hz to reduce a possible interference from high frequency activity such as extracellular spikes. Two minutes files were extracted from each recording and auto-correlograms and cross-correlograms were calculated using Spike2 software. To assess local gamma range synchronization, 2nd positive peak of auto-correlogram was measured for each region (CA3 or CA1). To determine gamma range synchronization between CA3 and CA1, the first positive peak of the cross-correlogram was measured.

### Proteomics

#### Mass spectrometry

Mass spectrometry was performed on vH samples collected from ~4- and ~9-weeks-old mice treated with or without DOX using an Orbitrap Fusion Lumos ETD mass spectrometer (Thermo) in data-dependent acquisition (DDA) mode. Peptides were separated on a C18 reversed-phase column with a linear gradient of acetonitrile in 0.1% formic acid at a flow rate of 300 nL/min. MS1 scans were acquired in the Orbitrap at a resolution of 120,000, covering a mass range of 375-1500 m/z. The top 3 most intense ions from each full scan were selected for MS/MS fragmentation using higher-energy collisional dissociation (HCD) at a normalized collision energy of 35%, with fragment ions detected in the Orbitrap at a resolution of 30,000. Dynamic exclusion was enabled to avoid repeated sequencing of the same peptides.

### Supplementary Figures

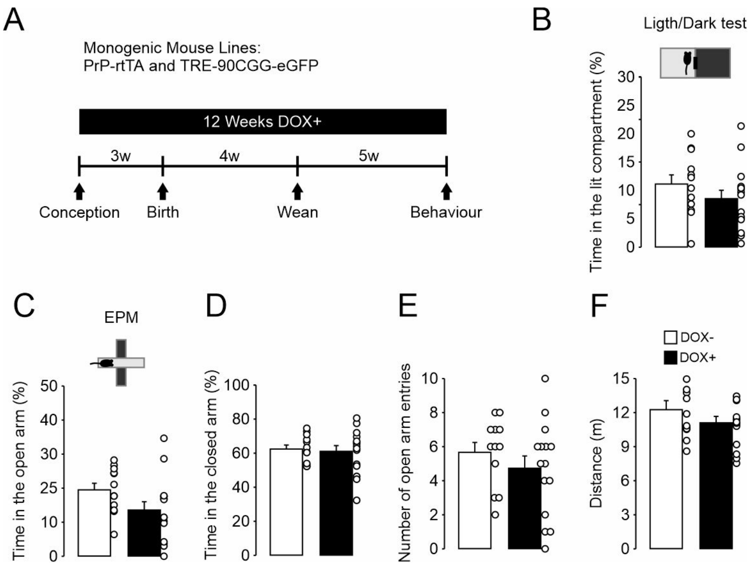


**Supplementary Figure 1 (Supplement to Figure 1) Doxycycline (DOX) treatment by itself does not cause changes in anxiety-like behaviour and locomotor activity in control Prp-rtTA and TRE-90CGG-eGFP monogenic mouse lines. (A)** Experimental timeline. (**B-F)** Monogenic mouse lines treated twelve weeks with DOX (DOX+: 15 mice) starting from embryonic development show similar anxiety-like behaviour to non-treated mice (DOX-: 12 mice) in **(B)** light-dark test ($t(25)=1.174$, $p=0.251$) and **(C-E)** elevated plus maze, with no changes in **(F)** locomotor activity (**C**: $t(25)=1.832$, $p=0.079$; **D**: $t(25)=0.307$, $p=0.761$; **E**: $t(25)=0.972$, $p=0.340$; **F**: $t(25)=1.221$, $p=0.234$). Statistical comparison for **b-f** Student´s two-tailed t-test. Data are presented as mean ± standard error of mean (SEM). Empty circles represent individual data points.

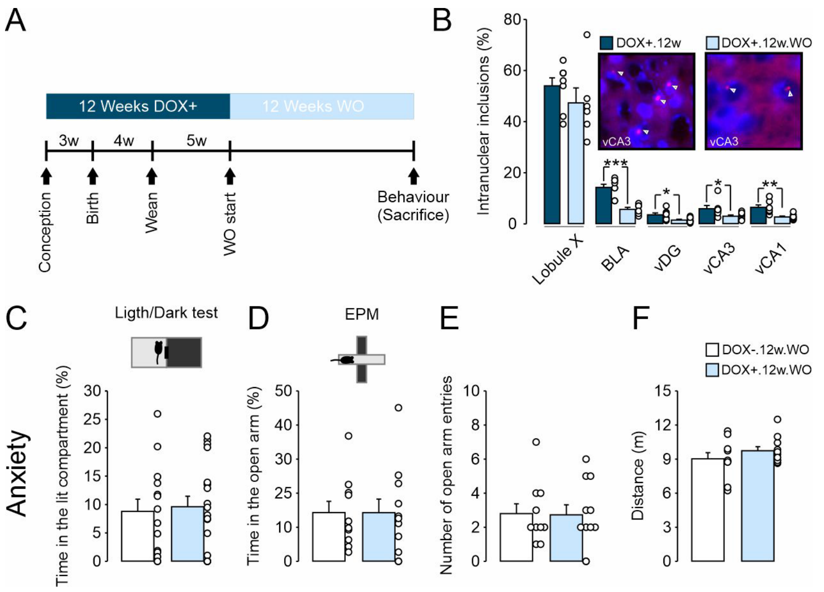


**Supplementary Figure 2 (Supplement to Figure 1) Cessation of doxycycline treatment for twelve weeks (DOX+.12w.WO) rescues anxiety-like behaviour and reduces the inclusion load in the limbic regions**. **(A)** Experimental timeline. **(B)** DOX+.12w.WO mice show lower levels of intranuclear inclusion in the basolateral amygdala (BLA, N=6 mice, t(12)=5.327, p<0.001) and ventral hippocampal (vH, N=8 mice) DG (U=8.5, p=0.021), CA3 (U=8, p=0.017) and CA1 (t(13)=3.932, p=0.002) subregions in comparison to DOX+.12w mice (BLA, N=8 mice; vH, N=7 mice). Note that the inclusion load in the cerebellum remains comparable to DOX+.12w mice (DOX+.12w: N=8 mice, DOX+.12w.WO: N=6 mice, t(12)=1.076, p=303). Representative intranuclear inclusions in the vCA3 of DOX+.12w.WO adult mice are highlighted with arrows. **(C-E)** DOX+.12w.WO mice show similar anxiety-like behaviour in **(C)** light-dark test (DOX-.12w.WO: N=14 mice, DOX+.12w.WO: N=18 mice, U=120, p=0.834) and **(D-E)** elevated plus maze (EPM; DOX-.12w.WO: N=11 mice, DOX+.12w.WO: N=12 mice) (**D**: t(19)=0.00526, p=0.996; **E**: t(19)=0.0727, p=0.931), and no differences in **(F)** locomotor activity (t(19)=-1.122, p=0.276). Statistical comparison for **B** (vDG, vCA3), **C**: Mann-Whitney Rank Sum Test. Statistical comparison for **B** (Lobule X, BLA, vCA1)**, D, E, F**: Student´s two-tailed t-test. Statistical differences are indicated via *p<0.05, **p<0.01 and ***p<0.001. Data are presented as mean ± standard error of mean (SEM). Empty circles represent individual data points.

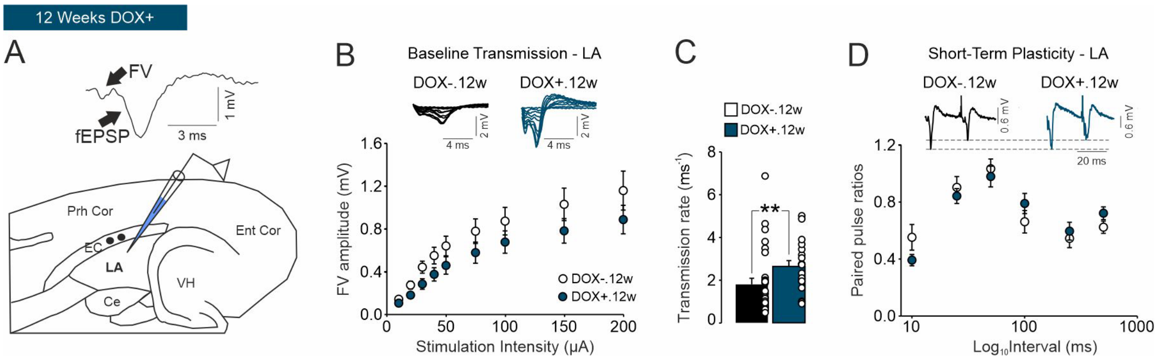


**Supplementary Figure 3 (Supplement to Figure 2) Increased baseline transmission in the lateral amygdala (LA) of young adult FXAND mice. (A)** Schematic presentation of recording configuration. Stimulation of external capsule (EC) induces extracellular field responses with fibre volleys (FV) and field excitatory postsynaptic excitatory potentials (fEPSP) in the LA. **(B)** FV amplitudes remain unchanged in DOX+.12w mice (DOX-.12w: N=10 mice, n=29 slices; DOX+.12w: N=7 mice, n=19 slices, (F(1, 46)=1.82, p=0.185). **(C)** Baseline transmission rate (fEPSP values normalized to FV amplitudes) is significantly increased (U=143, p=0.005) in DOX+.12w mice (N=10 mice, n=29 slices) compared to DOX-.12w mice (N=7 mice, n=19 slices). **(D)** Paired pulse responses are not altered in the LA of DOX+.12w mice (DOX-.12w: N=10 mice, n=30 slices; DOX+.12w: N=7 mice, n=22 slices, (F(1, 50)=0.000, p=0.997)). Statistical comparison for **B, D**: repeated measures two-way ANOVA followed by *posthoc* comparison using Holm-Sidak's multiple comparisons with Greenhouse–Geisser correction. Statistical comparison for **C**: Mann-Whitney Rank Sum Test. Statistical differences are indicated via **p<0.01. Data are presented as mean ± standard error of mean (SEM). Empty circles represent individual data points.

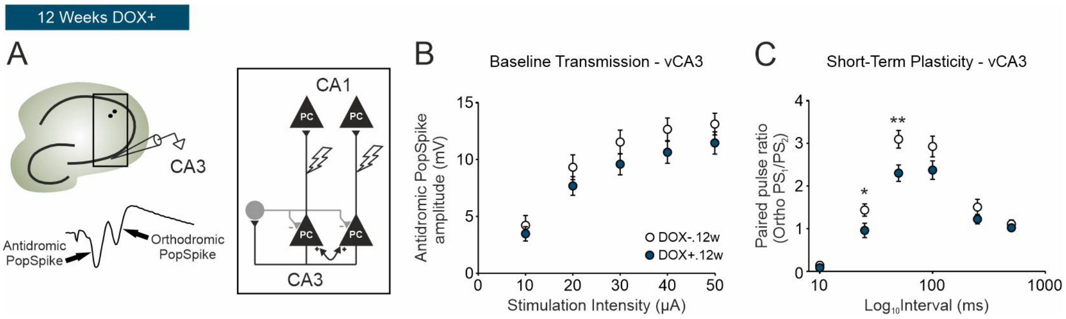


**Supplementary Figure 4 (Supplement to Figure 2) Altered short-term plasticity in the ventral CA3 (vCA3) of young adult FXAND mice. (A)** Schematic presentation of recording configuration. Antidromic stimulation of CA3 subregion (two black dots) induces an antidromic population spike and a following synaptically-mediated orthodromic population spike in the CA3. **(B)** Antidromic population spikes remain unchanged in DOX+.12w mice (DOX-.12w: N=5 mice, n=18 slices; DOX+.12w: N=5 mice, n=16 slices, $F(1, 32)=1.58$, $p=0.218$). **(C)** Paired pulse responses are reduced in the vCA3 of DOX+.12w mice (DOX-.12w: N=5 mice, n=18 slices; DOX+.12w: N=5 mice, n=16 slices, $F(1, 32)=10.9$, $p=0.0024$). Statistical comparison for **B, C**: repeated measures two-way ANOVA followed by *posthoc* comparison using Holm-Sidak's multiple comparisons with Greenhouse–Geisser correction. Statistical differences are indicated via *p<0.05 and **p<0.01. Data are presented as mean ± standard error of mean (SEM).

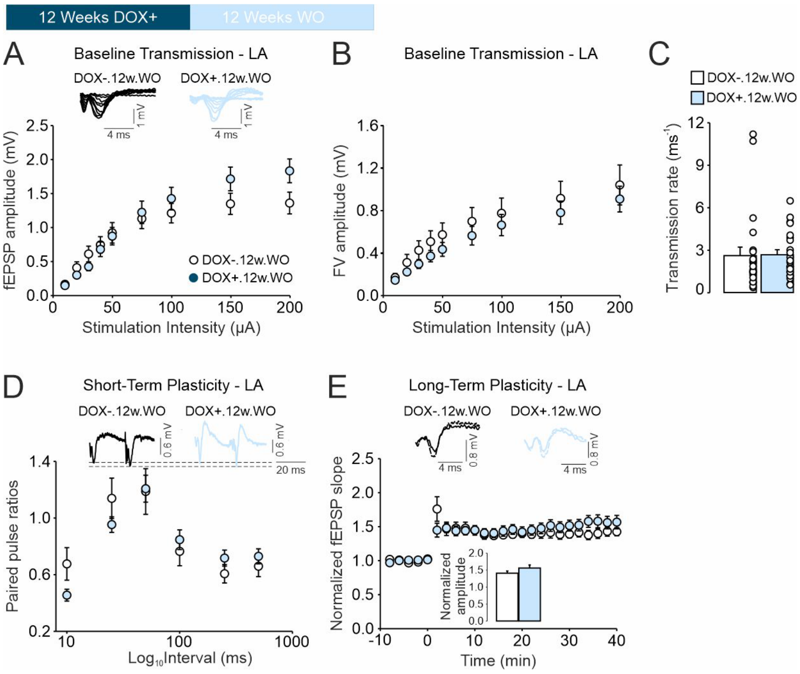


**Supplementary Figure 5 (Supplement to Figure 2) Cessation of doxycycline treatment for twelve weeks (DOX+.12w.WO) normalizes hyperexcitability in the lateral amygdala (LA).** **(A-B)** No main group effects are evident for excitability in the LA of DOX+.12w.WO mice evident by similar input-output curves for **(A)** fEPSP (DOX-.12w.WO: N=8 mice, n=24 slices; DOX+.12w.WO: N=8 mice, n=26 slices, F(1, 48)=0.291, p=0.592) and **(B)** FV amplitudes (DOX-.12w.WO: N=8 mice, n=23 slices; DOX+.12w.WO: N=8 mice, n=22 slices, F(1, 43)=0.749, p=0.392). **(C)** Baseline transmission rate (fEPSP values normalized to FV amplitudes) is not altered (U=205, p=0.281) in DOX+.12w.WO mice (N=8 mice, n=23 slices) compared to DOX-.12w mice (N=8 mice, n=22 slices). **(D)** Paired pulse responses are not altered in the LA of DOX+.12w.WO mice (DOX-.12w.WO: N=8 mice, n=24 slices; DOX+.12w.WO: N=8 mice, n=26 slices, F (1, 48)=0.0415, p=0.8394). **(E)** DOX+.12w.WO mice show no change in LA long-term potentiation (LTP) in comparison to DOX-.12w.WO mice (DOX-.12w.WO: N=6 mice, n=16 slices; DOX+.12w.WO: N=8 mice, n=14 slices, F (1, 30)=0.0997, p=0.754). Statistical comparison for **A, B, D, E**: two-way repeated measures ANOVA followed by posthoc comparison using Holm-Sidak's multiple comparisons with Greenhouse–Geisser correction. Statistical comparison for **C**: Mann-Whitney Rank Sum Test. Statistical comparison for **E** (Bar graph): Student´s two-tailed t-test. Data are presented as mean ± standard error of mean (SEM). Empty circles represent individual data points.

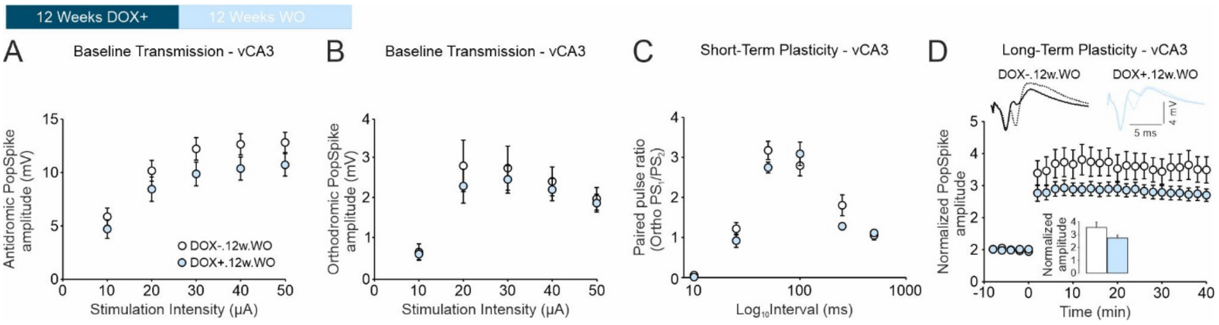


**Supplementary Figure 6 (Supplement to Figure 2) Cessation of doxycycline treatment for twelve weeks (DOX+.12w.WO) normalizes aberrant electrophysiological phenotype in the vCA3. (A)** Schematic presentation of recording configuration. **(A-B)** No main group effects are evident for excitability in the vCA3 of DOX+.12w.WO mice evident by similar input-output curves for **(A)** antidromic (DOX-.12w.WO: N=6 mice, n=14 slices; DOX+.12w.WO: N=8 mice, n=18 slices, F(1, 30)=1.84, p=0.1846) and **(B)** orthodromic population spikes (DOX-.12w.WO: N=6 mice, n=14 slices; DOX+.12w.WO: N=8 mice, n=18 slices, F(1, 30)=0.304, p=0.5855). **(C)** Paired pulse responses are not altered in the vCA3 of DOX+.12w.WO mice (DOX-.12w.WO: N=6 mice, n=14 slices; DOX+.12w.WO: N=8 mice, n=18 slices, F(1, 30)=1.21, p=0.2797). **(D)** DOX+.12w.WO mice show a non-significant trend towards reduction in long-term potentiation (LTP) in comparison to DOX-.12w.WO mice (DOX-.12w.WO: N=6 mice, n=10 slices; DOX+.12w.WO: N=8 mice, n=18 slices, F(1, 26)=3.65, p=0.0673). Statistical comparison for **A, B, C, D**: repeated measures two-way ANOVA followed by *posthoc* comparison using Holm-Sidak's multiple comparisons with Greenhouse–Geisser correction. Statistical comparison for **D** (Bar graph): Mann-Whitney Rank Sum Test. Data are presented as mean ± standard error of mean (SEM).

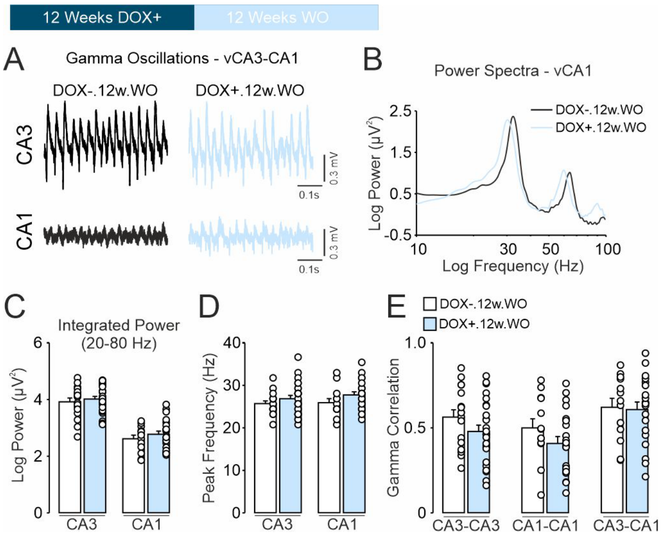


**Supplementary Figure 7 (Supplement to Figure 2) Cessation of doxycycline treatment for twelve weeks (DOX.+12w.WO) normalizes the enhanced gamma oscillations in the ventral hippocampus (vH). (A)** Representative traces of vCA3-CA1 and **(B)** vCA1 power spectra of cholinergic gamma oscillations illustrating unaltered gamma oscillations in the vCA3-CA1 axis of DOX+.12w mice (CA3, DOX-.12w.WO: N=6 mice, n=17 slices, DOX+.12w.WO: N=8 mice, n=25 slices; CA1: DOX-.12w.WO: N=6 mice, n=13 slices; DOX+.12w.WO: N=8 mice, n=22 slices). Summary data for gamma oscillations showing no statistically significant changes in **(C)** gamma power (20-80 Hz, CA3: $t(40)=-0.626$, $p=0.535$; CA1: $t(33)=-0.903$, $p=0.373$), **(D)** gamma peak frequency (CA3: $t(40)=-1.074$, $p=0.289$; CA1: $t(33)=-1.528$, $p=0.136$) and **(E)** gamma correlations in the vCA3-CA1 (CA3: $t(40)=1.479$, $p=0.147$; CA1: $t(32)=1.374$, $p=0.179$; CA3-CA1: $t(31)=0.191$, $p=0.850$). Statistical comparison for **C, D, E**: Student´s two-tailed t-test. Data are presented as mean ± standard error of mean (SEM). Empty circles represent individual data points.

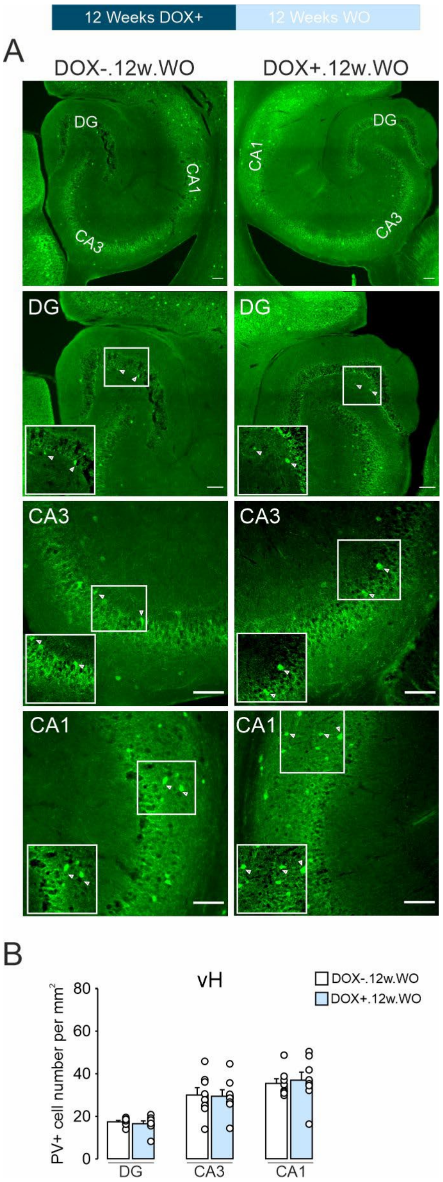


**Supplementary Figure 8 (Supplement to Figure 2) Cessation of doxycycline treatment for twelve weeks (DOX+12w.WO) normalizes the observed reduction in the parvalbumin+ (PV+) interneuron number in the ventral hippocampus (vH). (A)** Representative immunohistochemical stainings of PV+ interneurons in the vH of DOX+.12w.WO mice (N=8 mice per region) and DOX-.12w.WO mice (N=8 mice per region) and **(B)** summary data showing unaltered PV+ cell number in vH of DOX+.12w mice (DG: U=27, p=0.637; CA3: t(14)=0.126, p=0.931; CA1: t(14)=-0.352, p=0.730). Statistical comparison for **B** (vDG): Mann-Whitney Rank Sum Test. Statistical comparison for **B** (vCA3 and vCA1): Student´s two-tailed t-test. Data are presented as mean ± standard error of mean (SEM). Empty circles represent individual data points.

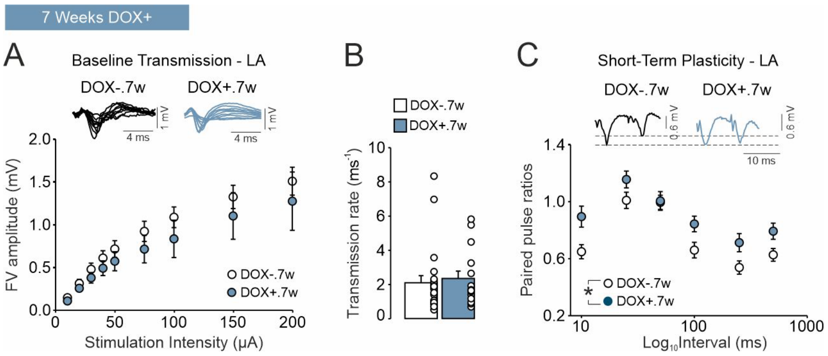


**Supplementary Figure 9 (Supplement to Figure 3) Altered short-term plasticity in the lateral amygdala (LA) of adolescent FXAND mice**. **(A)** FV amplitudes ($F(1, 35)=0.969$, $p=0.332$) and **(B)** baseline transmission rate ($U=143$, $p=0.506$) remain unchanged in DOX+.7w mice (N=10 mice, n=22 slices) compared to DOX-.7w mice (N=10 mice, n=15 slices). **(C)** Paired pulse responses are enhanced in the LA of DOX+.7w mice (DOX-.7w: N=10 mice, n=23 slices; DOX+.7w: N=10 mice, n=17 slices, $F(1, 38)=6.84$, $p=0.013$). Statistical comparison for **A, C**: repeated measures two-way ANOVA followed by *posthoc* comparison using Holm-Sidak's multiple comparisons with Greenhouse–Geisser correction. Statistical comparison for **B**: Mann-Whitney Rank Sum Test. Statistical differences are indicated via $^{*}p<0.05$ and $^{**}p<0.01$. Data are presented as mean ± standard error of mean (SEM). Empty circles represent individual data points.

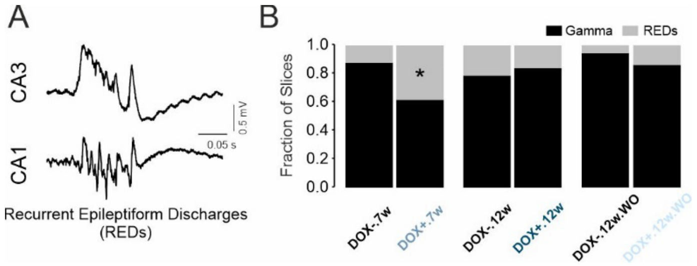


**Supplementary Figure 10 (Supplement to Figures 1-3) Propensity to generate recurrent epileptiform discharges (REDs) in the ventral hippocampus (vH) of adolescent FXAND mice is increased. (A)** Trace of a RED recorded in the vCA3-CA1. **(B)** Quantification of fraction of slices that generated REDs in DOX+.7w mice (~38%, 16 out of 42 slices) is significantly increased in comparison to DOX-.7w mice (~12%, 4 out of 33 slices). No difference is evident for altered RED generation in DOX+.12w (~16%, 3 out of 19 slices) vs. DOX-.12w mice (~21%, 4 out of 19 slices) or DOX+.12w.WO (~14%, 4 out of 29 slices) vs. DOX-.12w.WO mice (~5%, 1 out of 18 slices). Statistical comparison for **B:** Fisher´s Exact test. Data are presented as fraction of slices that exhibited either gamma oscillations or REDs.

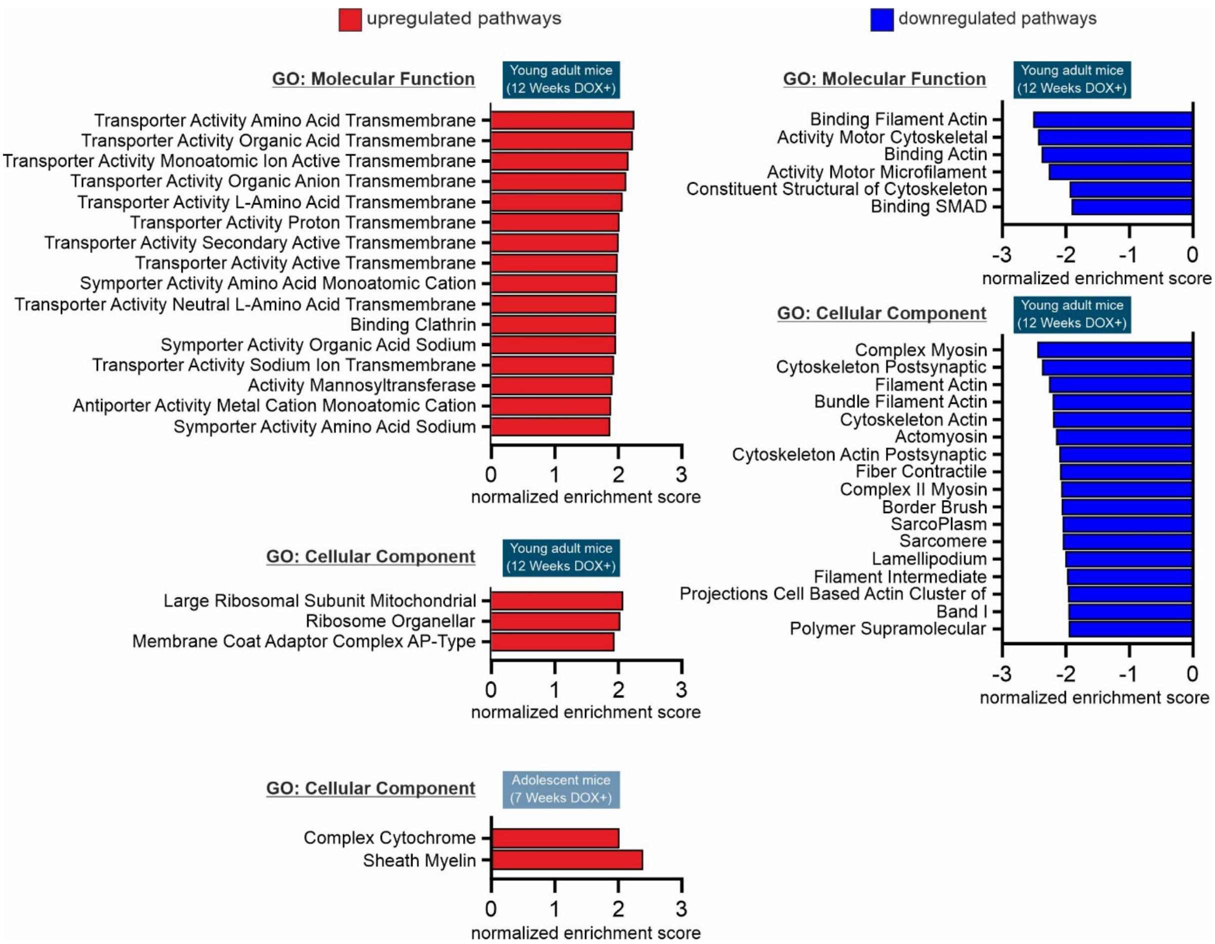


**Supplementary Figure 11 (Supplement to Figure 4) Upregulated and downregulated pathways identified in the ventral hippocampus of adolescent and young adult FXAND mice** with transgene activation using Gene ontology terms Molecular Function and Cellular Component. Note that profoundly more pathways are affected in the young adult mice in comparison to adolescent mice pointing to developmental stage and time dependent effects of transgene activation.
